# Evolutionary expansion of the corticospinal system is linked to dexterity in *Peromyscus* mice

**DOI:** 10.1101/2025.10.16.682851

**Authors:** Kelsey M. Tyssowski, Jeremy D. Cohen, Jian-Zhong Guo, Phoebe R. Richardson, Karen E. Cortina, Isobel H. Smith, Deshawn C. Eijogu, Adam W. Hantman, Hopi E. Hoekstra

## Abstract

While the central nervous system is perpetually reshaped by evolution, the principles governing how such changes promote ecologically relevant behaviors without breaking established functions are poorly understood. The expansion of neuron number is a potential mechanism by which the nervous system evolves to support changes in behavior without disrupting existing circuit function. Corticospinal neurons (CSNs) are a classic example: an expansion in the corticospinal system in the primate lineage has been hypothesized to underlie their exceptional dexterity. However, the role of CSN number in behavior has been difficult to assess due to the lack of a tractable model system. We compared two closely related subspecies of deer mice (*Peromyscus maniculatus*): forest mice, which evolved the ability to adeptly climb, presumably to support a semi-arboreal lifestyle, and prairie mice, which are less proficient climbers. We find that forest mice have about two-fold larger corticospinal tracts (CSTs) driven by an increase in CSN number in secondary motor and sensory cortical areas (M2 and S2). Furthermore, forest mice display greater manual dexterity than their prairie counterparts in a reach-to-grasp task, consistent with the idea that an increase in CSN number supports more dexterous behavior. High-throughput neural recordings during this task revealed a difference in the timing of neural activity between forest and prairie mice, specifically in M2: in forest mice, the peak of activity was shifted towards the grasping phase of the behavior. Forest mice also outperform their prairie counterparts on an ecologically relevant climbing task, where they spend more time upright crossing a thin rod, move faster, and right themselves more quickly when they fall, suggesting a general difference in motor dexterity not restricted to hand use. Finally, we use F2 hybrid animals to show that CST size is correlated with climbing dexterity, providing support for the long-standing hypothesis that corticospinal system expansion supports the evolution of dexterity. Together, our work establishes the forest-prairie deer mouse system as a model to investigate the role of neuron number expansion, and CSNs in particular, in dexterous movement.

## Introduction

The vast neuroanatomical variation across extant species suggests that evolution has tuned neuroanatomy to support behavioral adaptations to distinct ecological niches. Neuroanatomical variation falls into two non-mutually exclusive broad categories. First, circuit connectivity varies across taxa^1–6^, which may impact circuit activity^7^. Additionally, evolution has also changed neuron number^8–11^, which, over long time scales, manifests as generation of new brain regions and expansion of existing ones^12^. A prime example of such expansion is that of cortex in the primate lineage, which is thought to confer increased dexterity and cognition^13,14^. However, how the evolutionary expansion of specific cortical cell populations mediates this behavioral adaptation remains largely unknown.

One such cortical cell population that has undergone evolutionary expansion is corticospinal neurons (CSNs), which project from cortex to spinal cord. The expansion of the corticospinal system, both through changes in the number of CSNs and through changes in their synaptic partners in the spinal cord, has been hypothesized to underlie the evolution of dexterity^15–24^. Consistent with this idea, experiments across taxa demonstrate that manipulating CSNs or the corticospinal tract (i.e., axons of CSNs; CST) impairs both fine-motor control of the hand25–30, as well as complex locomotion^31,32^. Thus, CSNs have a clear role in dexterity, which has been defined as “finding a motor solution for any external situation”^33^. However, the role of CSN *number* in mediating dexterity has been difficult to test directly due to the lack of suitable evolutionary model systems that are both experimentally tractable and minimally evolutionarily diverged.

To overcome this challenge, we take advantage of two closely related subspecies of deer mice (*Peromyscus maniculatus*) that have evolved differences in dexterous movement. Following the last glacial retreat (∼10k years ago), North American deer mice living in prairie habitats migrated into forests where they evolved dexterous climbing behavior, presumably to support a semi-arboreal lifestyle, whereas populations living in prairies did not evolve this behavior^34–42^. This evolution occurred multiple times across North America, resulting in at least four convergently evolved forest-mouse subspecies (Figure 1A)^37^. These subspecies are not only closely related but also interfertile, enabling the generation of hybrids to aid in identifying the genetic basis of evolved variation between subspecies^40,43^. Compared to their prairie counterparts, forest mice have evolved shared morphological phenotypes, including longer tails and larger hind feet, that could also impact their behavior^36–38,40,41,43,44^. However, these morphological changes are minimal compared to, for example, the difference between a rodent and primate hand, making it easier to account for them experimentally. In addition, behavioral studies of wild-caught and lab-raised forest and prairie mice reveal a dramatic behavioral difference: forest mice outperform prairie mice on tests of both vertical and horizontal climbing^39–42^. Given the known role of the corticospinal system in dexterous behavior and the dramatic difference in dexterity between these subspecies, we hypothesized that the evolutionary expansion of the corticospinal system mediates evolution of dexterity in deer mice.

**Figure 1:**
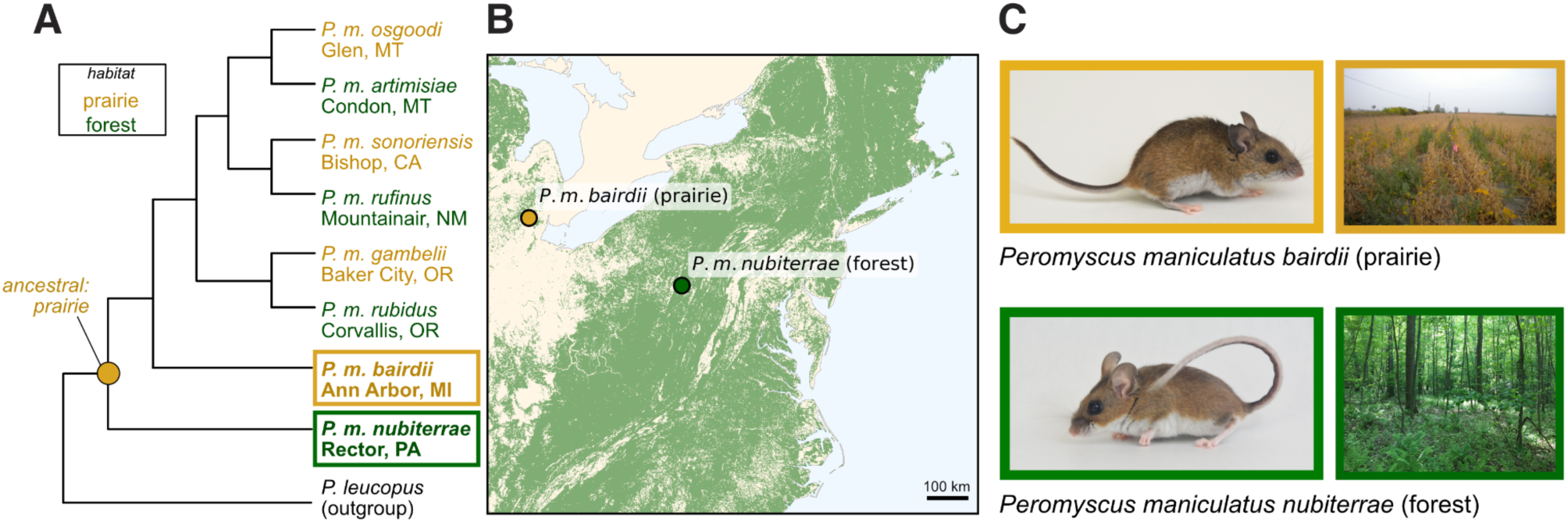
Forest and prairie deer mouse subspecies. A. Cladogram showing repeated evolution of forest and prairie deer mouse subspecies. The subspecies used in this study are outlined. B. Terrain map showing the trapping locations of the founders used to establish the mouse colonies used in this study. Green shading represents forests, specifically regions with greater than 30% tree canopy cover according to the United States Geological Survey National Land Cover Database. C. Photos of a representative prairie and forest mouse from the laboratory colony, and the respective habitats at the trapping locations from which the founders of those colonies were collected, as shown in B.

## Results

### Forest mice have larger corticospinal tracts compared to prairie mice

We focus on one forest and one prairie mouse subspecies, which are recently diverged (by less than 1 million years^45^) and exist as laboratory colonies: *P. m. nubiterrae* (forest) and *P. m. bairdii* (prairie)^36^ (Figure 1B-C). To determine whether CST size differs between subspecies, we labeled the CST using an antibody against protein kinase C gamma (PKCγ)^46,47^ in cleared, whole-mount spinal cords. Forest mouse spinal cords showed a greater PKCγ-positive CST volume, as assessed by lightsheet imaging, compared to their prairie counterparts (Figures 2A-E). This difference was particularly striking in the cervical spinal cord, which controls the forelimbs, where forest mice showed approximately 2-fold greater CST volume than prairie mice (Figures 2A,C), though we also observed larger thoracic/lumbar CSTs (Figure 2D). While we found a 1.4-fold difference between subspecies in total spinal cord volume (Figure S1A), this difference was entirely due to differences in lumbar spinal cord volume, with forest mice having 1.8-fold larger lumbar spinal cords, but no difference in cervical spinal cord volume (Figure S1D-G). Nonetheless, we observe significant differences in CST size even when normalizing for total spinal cord volume (Figure S1B). The particularly large CST expansion in the cervical spinal cord of forest mice suggests that as axons travel down the spinal cord, an increased proportion of axons branch off and innervate the grey matter of the cervical spinal cord in forest mice compared to prairie mice. Consistent with this conclusion, we observed a larger fraction of the total PKCγ volume in cervical spinal cord in forest mice compared to prairie mice (Figure 2E). We did not find differences in CST length (Figure S1C), suggesting that the CST targets the same spinal segments in both subspecies. Together, these data support the conclusion that the CST is larger in forest mice compared to prairie mice, consistent with our hypothesis that CST expansion may underlie the better dexterity of forest mice.

**Figure 2:**
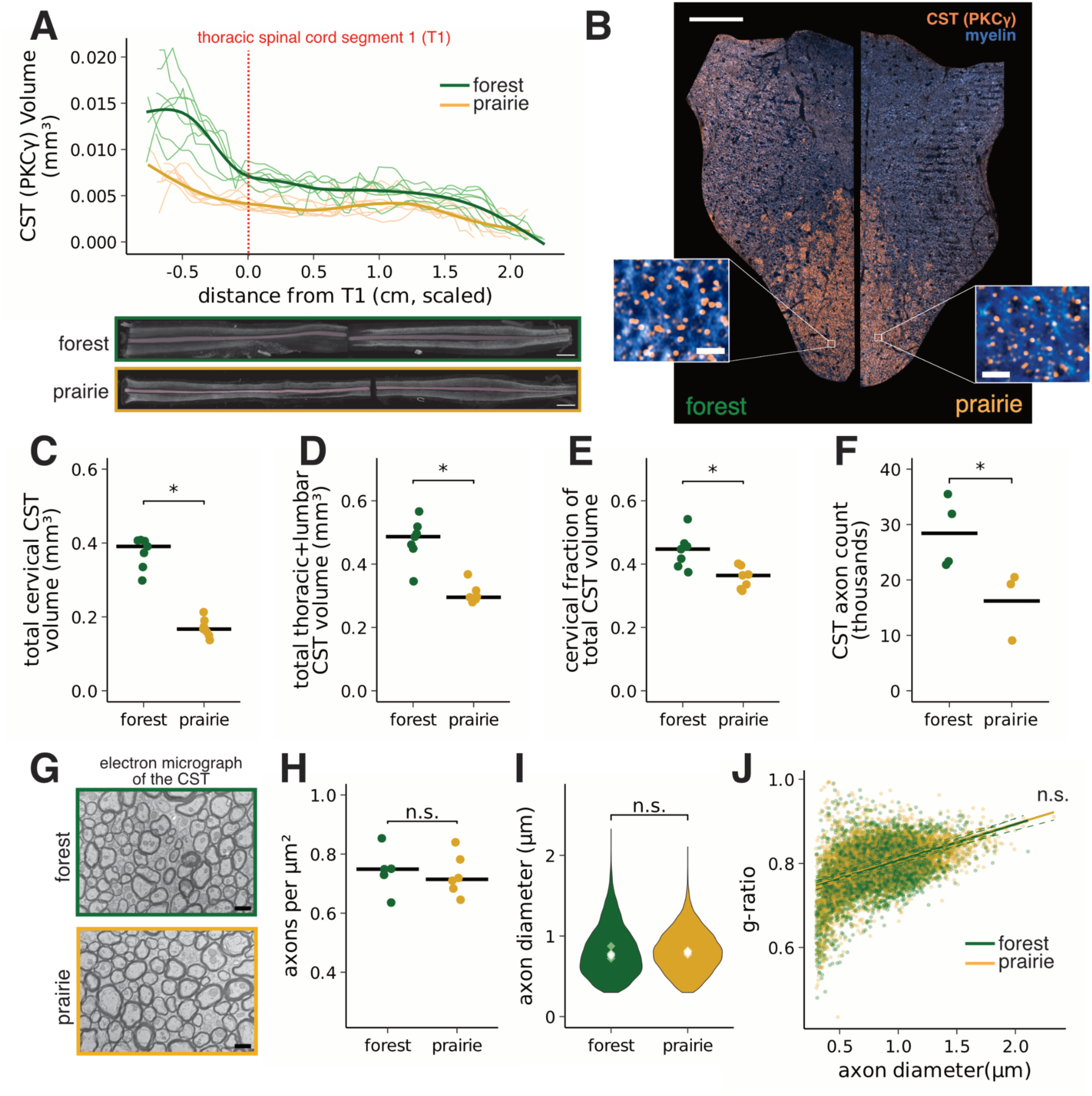
Forest mice have a larger corticospinal tracts (CSTs) due to an increase in axon number. A. Comparison of PKCγ+ corticospinal tract size between forest and prairie mice in whole-mount, cleared spinal cords. Top: Lines represent PKCγ+ corticospinal volume binned at 140µm and smoothed. Thin lines represent individual animals, and thick lines are a fitted regression for each subspecies. T1 marks the beginning of the thoracic spinal cord. n=7 animals of each subspecies. *Bottom:* Representative projections along the dorsal/ventral axis of 3D images of forest and prairie mouse spinal cords with PKCγ staining (cervical and lumbar segments imaged separately). The corticospinal tract is highlighted in pink. The other signal is PKCγ+ interneurons. Scale bar=1mm. B. Representative images of coronally sectioned hemisections of forest and prairie mouse cervical segment three dorsal funiculi stained with FluoroMyelin dye to label myelin and immunostained for PKCγ. Scale bar for full-sized image=100µm. Scale bar for zoomed insets=2µm. C. Quantification of data shown in A. Points represent values for individual animals. Black lines represent the median of each subspecies. *p=0.00073, linear regression comparing cervical CST volume between subspecies, accounting for sex, age, staining batch, and total spinal cord volume (β=0.19, 95% CI: 0.106–0.270, t₈=5.29). D. Quantification of data shown in A. Points represent values for individual animals. Black lines represent the median of each subspecies. *p=0.00021, linear model comparing thoracic+lumbar CST volume between subspecies, accounting for sex, age, staining batch, and total spinal cord volume (β=9.68, 95% CI: 6.19–13.17, t₈=6.39). E. Quantification of data shown in A. Points represent values for individual animals. Black lines represent the median of each subspecies. *p=0.007, two-sided Wilcoxon rank-sum test (median [IQR]: prairie, 0.364 [0.328–0.381]; forest, 0.447 [0.405–0.460]; Z=-2.61, r=0.70). F. Quantification of the number of PKCγ+ axons in a hemisection of the dorsal funiculus (prepared as in B) from n=4 forest and n=3 prairie mice. Points represent values for individual animals. Black lines represent the mean of each subspecies. *p=0.034, two-sided Wilcoxon rank-sum test (median [IQR]: prairie, 19138 [14210–19674]; forest, 27807 [23481–32743]; Z=-2.12, r=0.80). G. Representative electron micrographs of the ventral part of forest and prairie coronally sectioned dorsal funiculi. Scale bar = 1µm. H. Quantification of axon density in electron micrographs from n=6 animals of each subspecies. Points represent values for individual animals. Black lines represent the median of each subspecies. p=0.748, two-sided Wilcoxon rank-sum test (median [IQR]: prairie, 0.710 [0.683–0.723], n=6; forest, 0.732 [0.648–0.759], n=6; Z=-0.40, r=0.12). I. Quantification of axon diameter in electron micrographs from n = 6 animals of each subspecies. Violin plots represent the distribution of axon diameters of all axons measured. White diamonds represent the medians of each animal. p=0.132, two-sided Wilcoxon rank-sum test (median [IQR]: prairie, 0.803 [0.788–0.808], n=6; forest, 0.768 [0.756–0.780], n=6; Z=1.60, r=0.46). J. Quantification of g-ratio (axon diameter/axon+myelin diameter) from n=6 animals of each subspecies. Points represent g-ratios of individual axons measured. Dashed lines are the trend lines for individual animals, and solid lines are the trend lines for each subspecies. p=0.70, two-sided Wilcoxon rank-sum test (median [IQR]: prairie, 0.794 [0.791–0.799]; forest, 0.796 [0.792–0.806]; Z=- 0.39, r=0.11).

The difference we observe in CST volume could be due to an increase in the number of CST axons or to differences in the density or morphology of individual axons. To distinguish between these possibilities, we counted axons stained with FluoroMyelin, a fluorescent myelin stain, within the PKCγ-positive CST region in coronal sections of the dorsal funiculus of the cervical spinal cord (Figure 2B). We found that the number of axons in the CSTs of forest mice was about 2-fold greater than that of prairie mice, consistent with the increase in CST volume being due primarily to an increase in axon number (Figure 2F). To assess the ultrastructure of CST axons between subspecies, we performed electron microscopy on the ventral region of the dorsal funiculus, where the CST resides (Figure 2G). We found no difference between subspecies in axon density (Figure 2H), axon diameter (Figure 2I), or myelination, as measured by the g-ratio (i.e., the ratio of the total axon diameter to the axon diameter inside the myelin; Figure 2J). These data support the idea that the larger CST size in forest mice is primarily driven by an increase in the number of axons, rather than differences in axon density or morphology. Although this does not entirely rule out the possibility that more axons branch before exiting the CST in forest mice, the most parsimonious explanation for this increase in axon number is a corresponding increase in CSN number, which we next sought to test directly.

### Forest mice have more corticospinal neurons in secondary motor and sensory cortex than prairie mice

In rodents, CSNs reside in several cortical areas: primary motor cortex (M1, also called caudal forelimb area; CFA), secondary motor cortex (M2, also called rostral forelimb area; RFA), primary sensory cortex (S1), and secondary sensory cortex (S2)^48^. To determine whether the increase in CST size results from an expansion of a specific spatially defined subpopulation of CSNs, we labeled CSNs by injecting rAAV2-retro^49^ expressing nuclear-tagged GFP into the cervical spinal cord grey matter. We injected the grey matter of one spinal segment between C5-C8 to target cervical spinal cord (Figures 3A, S2A), where we expected, from our CST volume measurements, to observe a difference in CSN grey matter innervation between subspecies. We then quantified nuclei throughout the brain in light-sheet images of cleared brains and assigned them to cortical regions (or the rest of the brain) using a custom pipeline. To control for differences in viral labeling, we normalized our counts to the total number of labeled nuclei per brain, which did not differ between subspecies, nor did total brain volume (Figure S2B-C). We observed more CSNs in M2 and S2 in forest compared to prairie mice, but a similar number in M1 and S1 (Figure 3B-D). These data indicate that the increase in CST size is due to an increase in CSN number, specifically in secondary, but not primary, motor and sensory cortex.

**Figure 3:**
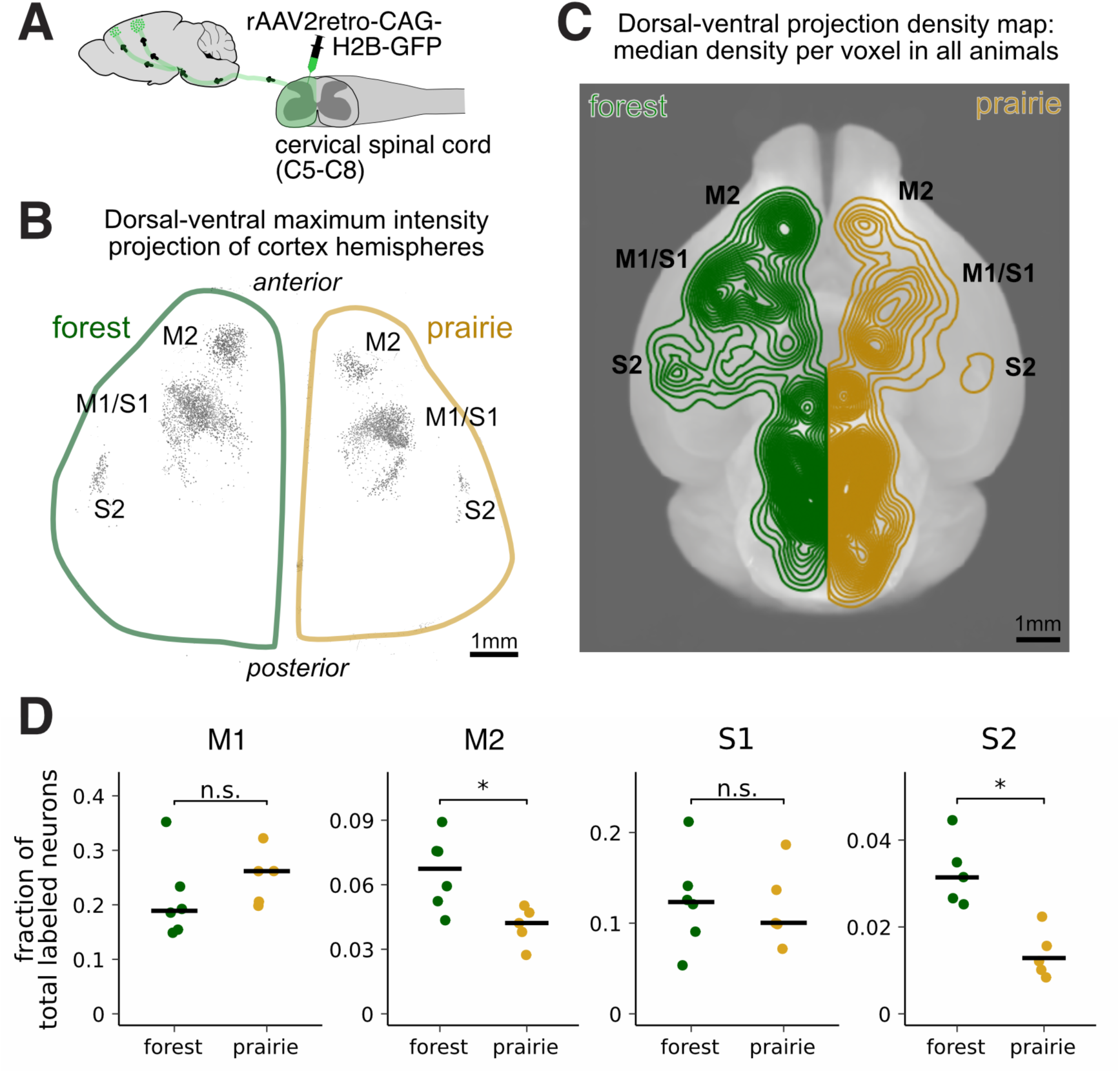
Forest mice have more corticospinal tract neurons specifically in secondary motor and sensory regions (M2 and S2) compared to prairie mice. A. Schematic of the injection. Retrograde AAV carrying a nuclear-tagged-GFP payload was injected into segments C5-C8 of the spinal cord. We analyzed brains with tagged nuclei. B. One representative brain from each subspecies showing a maximum intensity projection across the dorsal/ventral axis of the signal from labeled neuronal nuclei in contralateral cortex. Cortical areas labeled are primary motor (M1) and sensory (S1) cortex, and secondary motor (M2) and sensory (S2) cortex. The signal from M1 is in caudal forelimb area (CFA), and from M2 is in rostral forelimb area (RFA). C. Density plot of the median fraction of nuclei labeled across the contralateral (to the injection) side of the brain. Data are shown as a projection along the dorsal/ventral axis, with cortical areas labeled (as in B). Data from n = 6 forest mouse brains, n = 5 prairie mouse brains. D. Quantification of the fraction of total labeled neurons in each brain region labeled in B/C. *FDR-adjusted q<0.05, based on two-sided Wilcoxon rank-sum tests (M1: Z=1.46, r=0.44, q=0.19; M2: Z=-2.37, r=0.72, q=0.035; S1: Z=-0.18, r=0.06, q=0.86; S2: Z=-2.61, r= 0.83, q=0.035).

CSNs are a subtype of neurons in layer V of cortex. We therefore considered the possibility that the CSN number expansion is due to a broader expansion of layer V or of sensorimotor cortex as a whole. We performed plate-based single-nucleus RNA sequencing (snRNA-seq) of frontal cortex, including all regions where CSNs reside, in five forest and five prairie mice, assessing a total of 107,000 cortical nuclei. We integrated data from individual nuclei between datasets to compare between subspecies, clustered nuclei by gene expression, and aligned our data to the BICNN atlas^50^ to identify cell types (see Methods). We observed no difference in the total number of neurons (or other cell types) between subspecies, indicating that forest mice do not have a general increase in neuron number across frontal cortex (Figure S3A-D). At the level of neuron subtypes, we observed only a small but significant increase in the number of L6b neurons in prairie mice (Figure S3E-G). Notably, we were able to clearly distinguish layer V extratelencephalic (L5ET) neurons (a subclass that includes CSNs, as well as corticopontine, corticotectal, corticorubral neurons, and others^50,51^). Still, we observed no difference in L5ET neuron abundance between subspecies. We note that the failure to observe a difference in abundance in L5ET neurons is not surprising given that CSNs are expected to make up only a small fraction of L5ET neurons, and suggests that the increase in CSN number is specific to CSNs rather than a side effect of increasing the size of layer V as a whole. Consistent with previous work suggesting that virally labeling CSNs is necessary to identify them in snRNA-seq data^52,53^, we were unable to identify a clear CSN cluster among L5ET neurons (Figure S3H-K). However, we asked whether we could identify putative markers of the expanded CSN population through differential gene expression. Pseudo-bulk differential gene expression analysis between forest and prairie mice L5ET neurons revealed a significant enrichment in known CSN marker genes in the genes upregulated in forest compared to prairie mice (Figure S1L) and identified several known CSN markers that could mark the expanded CSN population. Therefore, CSN expansion in forest mice is not likely a side effect of a general increase in neuron number or in L5ET neurons but is specific to a subset of CSNs that may be distinguishable by specific CSN marker genes.

### Forest mice display more dexterous hand use than prairie mice

Given the role of cortex, and CSNs more specifically, in dexterity, we next asked to what extent forest and prairie mice differ in dexterous movement. While forest and prairie mice differ in their ability to climb^34,35,54,36–42^, it is unclear whether they also differ in other, more common, measures of dexterity. Because many studies across taxa have demonstrated the role of the CST in a reach-to-grasp task^25–28^, we assessed forest and prairie mice in this behavioral paradigm, hypothesizing that forest mice would be more successful at this task given their expanded CSTs. We measured the success of animals reaching for a food pellet through a slit over 6 days of training. We found that forest mice were more successful at this task both at the beginning and after training compared to their prairie counterparts (Figure 4A), indicating that forest mice are more dexterous than prairie mice, consistent with their greater number of CSNs.

**Figure 4:**
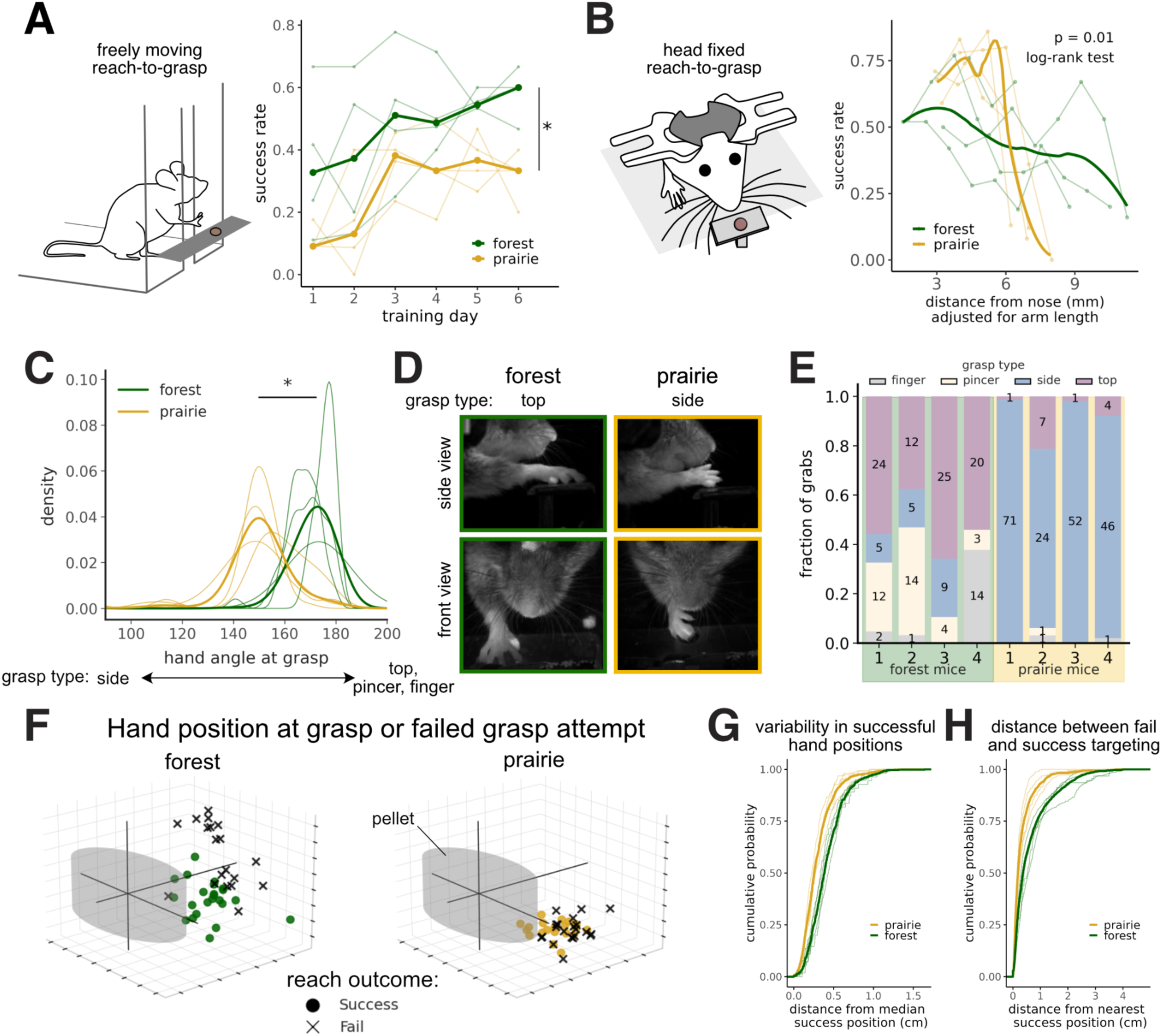
Forest mice show enhanced dexterity in a reach-to-grasp task compared to prairie mice. A. Comparison of forest and prairie mouse performance in a freely moving pellet-grasping task over 6 days of training. Thin lines represent individual animals (n=4 of each subspecies), thick lines represent the subspecies median. Forest mice successfully grasp the pellet at a higher rate, *p=0.018, F=10.4, repeated measures ANOVA on the difference between subspecies. B. Comparison of forest and prairie mouse performance in a head-fixed pellet-grasping task. Pellet position is adjusted for arm length. Thin lines represent individual animals (n=4 of each subspecies), thick lines represent the subspecies median. Forest mice reach further than prairie mice before completely failing (p=0.01, χ²=7.3, log-rank test). C. Distribution of the angle of the hand relative to the platform in forest and prairie mice during successful reach attempts at their most successful pellet position, with 180 degrees being parallel to the platform (top grasp type). Lighter lines show distributions of individual animals, darker lines show the distributions of the subspecies. Forest mouse hand positions are more parallel to the platform (top-down reach) at the time of grasp, *p=0.021, two-sided Wilcoxon rank-sum test on per-animal median hand angles (median [IQR]: prairie, 149.45 [148.73–151.22]; forest, 171.61 [169.57–174.05]; Z=-2.31, r=0.82; n=4 animals of each subspecies). D. Examples of images of forest and prairie mouse typical grasp types at the time of the start of the grasp. E. Quantification of the frequency of four qualitative grasp types in forest and prairie mice. Data quantified are from each animal’s preferred (i.e., most successful) pellet position. Forest mice have a greater diversity of grasp types. Grasp types are defined as follows: top = an approach from the top that uses all fingers to grasp the pellet; side = a scooping approach from the side; pincer = a grasp in which the pellet is grasped resting on the thumb-like digit one with another finger; and finger = when the pellet is grasped in between two fingers. F. Example data plotting the 3D tracked hand position, i.e., center of the hand, at the time of grasp (or grasp attempt in the case of a failed trial). Each plot shows the data from one animal on one day at their preferred pellet position. G. Cumulative distribution plot showing, for each successful reach attempt, the variability of target positions, i.e., the distance of the hand at grasp from the median successful hand position. Light lines show individual animals, dark lines show subspecies distributions. Forest mice have a wider range of success positions than prairie mice (p=0.029, two-sided Wilcoxon rank-sum test comparing per-animal median absolute deviations; median [IQR]: prairie, 0.264 [0.250–0.273]; forest, 0.369 [0.350–0.398]; Z=-2.31, r=0.82; n=4 animals of each subspecies). H. Cumulative distribution plot showing, for each failed reach attempt, the closest the hand comes to a successful grasp position. Light lines show individual animals, dark lines show subspecies distributions. Forest mice are further from a successful position when they fail, suggesting forest mice fail more often due to targeting their hand to the wrong position, whereas prairie mice fail at grasping even if their hand is targeted well (p= .029, two-sided Wilcoxon rank-sum test comparing per-animal fail-success distances; median [IQR]: prairie, 0.176 [0.152–0.206]; forest, 0.469 [0.376–0.541]; Z=-2.31, r=0.82; n=4 animals per subspecies),

We next sought to understand the behavioral mechanisms underlying this difference in success. We used a head-fixed version of the reach-to-grasp task to afford us better control over task parameters. In this version of the task, we were able to train prairie mice to achieve a success rate equivalent to that of forest mice (Figure 4B, Figure S4A). However, we found that prairie mice were only able to successfully grasp pellets close to their noses, whereas they failed at more distant positions. In contrast, when we presented forest mice with pellets progressively further from their noses, they were able to successfully grasp the pellet at all assessed distances (Figure 4B). Importantly, this difference in performance cannot be explained by differences in limb length. While the median forelimb length of forest mice is about 3mm longer than that of prairie mice^55^ (consistent with measurements in our sample, Figure S4B), the difference in the distance from the nose at which they can successfully grasp a pellet is approximately 7mm (13.8+/-1.6mm in forest mice, 7.1+/-0.8mm in prairie mice). This large difference in distance at which mice can succeed indicates that differences in arm length cannot entirely account for differences in the ability to grasp more distant pellets, and that forest mice have greater flexibility in the pellet positions they can successfully reach compared to their prairie counterparts.

Previous reports in both rats and monkeys demonstrate that lesioning the CST alters the dexterous grasping part of the reach-to-grasp behavior. Specifically, animals transition from using dexterous grasping to using a scooping movement^27,28^. We therefore hypothesized that forest animals may be more likely to use a dexterous, top-down grasping movement, whereas prairie mice use a less dexterous scooping movement. Indeed, we found that forest mice approach the pellet with their hands parallel to the platform. In contrast, prairie mice approach the pellet with their hands at a more perpendicular angle to the platform to execute a scooping grasp (Figure 4C-D, Videos S1-2). We note that the grasping angle is not dependent on pellet position (Figure S4D-E). We next asked whether forest and prairie mice use qualitatively different grasping strategies. We divided grasps into four categories: top, describing an approach from the top that uses all fingers to grasp the pellet; side, describing a scooping approach from the side; pincer, which describes a grasp in which the pellet rests on the thumb-like digit one with another finger; and finger, which describes when the pellet is in between two fingers (Figure S4C, Videos S3-6). When we compared grasps at each mouse’s most successful position, we found that forest mice use a wider variety of grasp types compared to prairie mice, which, with a small number of exceptions, exclusively use the side grasp (Figure 4E), suggesting that forest mice are more dexterous in the grasping part of the behavior, consistent with our hypothesis that corticospinal expansion supports greater dexterity.

We reasoned that the more varied grasping strategy of forest mice may enable them to successfully grasp the pellet even if they inconsistently target their hands to the pellet location. In contrast, because prairie mice may only be able to grasp the pellet in one way, they need to be able to direct their hands to the optimal position to perform that grasp. To determine the variability in the position of the hand during successful grasps, we calculated the distance between the median successful hand position at grasp and each successful grasp position. We found that forest mice displayed greater variability in successful grasp positions (i.e., higher median absolute deviations), consistent with the idea that their increased flexibility in grasp types enables more flexibility in pellet targeting (Figure 4F-G). Consistent with this finding, we also observed that forest mice had greater variability not only in their targeting but also in the overall trajectory to the pellet (Figure S4I-J). We note that forest and prairie mice significantly differ in their pellet targeting and reach trajectory even when accounting for pellet position (Figure S4G,J). We considered whether the difference in targeting might be due to differences in hand morphology, as forest mice have slightly longer fingers than prairie mice, which could allow them to grasp the pellet from a larger surrounding radius. We therefore measured how far each successful grasp position was from the pellet itself, but found this was not different between subspecies (Figure S4F). Furthermore, a linear mixed effects model accounting for this hand-pellet distance showed a significant impact of subspecies (p = 0.0003; see Figure S4G, Methods), supporting our conclusion that forest mice use a wider range of grasp strategies that is not merely a result of their differing morphology.

To investigate the role of dexterity in success on the task, we assessed how forest and prairie mice fail. We reasoned that if forest mice indeed have a better ability to grasp the pellet than prairie mice, a larger fraction of prairie mouse failures would be failures in grasping, whereas forest mouse failures would be in targeting. To address this, for each failed reach, we assessed the nearest point in the reach trajectory to the grasp position of a successful reach-to-grasp (fail-success distance). We defined a reach that never comes close to a successful reach position as a failure in targeting, while a reach in which the mouse’s hand arrives at a location where the animal has made a successful grasp is more likely to be an error in grasping itself. A comparison of the distribution of fail-success distances between forest and prairie mice supported our hypothesis that while forest mice are more likely to fail by targeting poorly, prairie mice often fail despite targeting their hands to a position where they were previously successful (Figure 4F,H). In support of the idea that a difference in grasping dexterity might impact the ability of prairie mice to successfully grasp pellets far from their noses, we find that at far pellet distances, prairie mouse failures are more often targeting failures (Figure S4H).

This difference between subspecies suggests that, at greater distances, where targeting becomes harder, grasping dexterity is more critical for success. Together, these data further support the idea that forest mice are more dexterous in their reach-to-grasp behavior. While prairie mice can learn to be successful at the task in a confined set of circumstances in which they can easily, consistently, and precisely target the ideal grasp location, forest mice, presumably due to their increased manual dexterity, do not need to achieve precise targeting to achieve similar levels of success.

### Neural dynamics in forest mouse M2 differ from those of prairie mouse M2

Given the role of cortex in dexterous behavior and the observed differences in cortical neuroanatomy, we asked whether differences in CSN number between forest and prairie mice might impact neural dynamics. Because forest and prairie mice differ in the extent to which grasping versus targeting contribute to their success in the reach-to-grasp task, we hypothesized that this difference in behavior might be reflected in motor cortex activity. Using Neuropixels probes^56^, we focused on motor areas with most evidence of involvement in the task^23,26,29,57–60^, recording from M1 (a.k.a. CFA, where we observed no CSN number differences between subspecies) and M2 (a.k.a. RFA, where we observed differences between subspecies) in trained, head-fixed animals during reaching behavior. We recorded from the same animals over 3-5 sessions and validated probe placement by comparing it to our retrogradely labeled brains (Figure S5A). We observed robust neural signals aligned with the behavior, similar to that observed in lab mice in the same conditions^57^. We aligned neural signals during successful reaches to the time of the grasp. Interestingly, we observe a difference in the timing of the signal between forest and prairie mice in M2, but not in M1 (Figure 5A-C, S5B-E). In prairie mice, the neural signal in M2 peaks before the time of the grasp and before the signal in M1, as expected from recordings in lab mice^58^. In contrast, in forest mice, the M2 signal is later, more aligned with the timing of the grasp and more similar to the M1 signal. Notably, this is only true for positively modulated neurons (i.e., those that increase with behavior), but not for negatively modulated neurons, which show similar alignment in activity between forest and prairie animals (Figure 5A-B, S5B-E). These findings indicate that neural dynamics in cortex differ between forest and prairie mice and are consistent with our behavioral results. Furthermore, the fact that we observe a difference specifically in M2 suggests that the difference in CSN number, which we observe in M2 but not M1, may play a role in this difference in dynamics.

**Figure 5:**
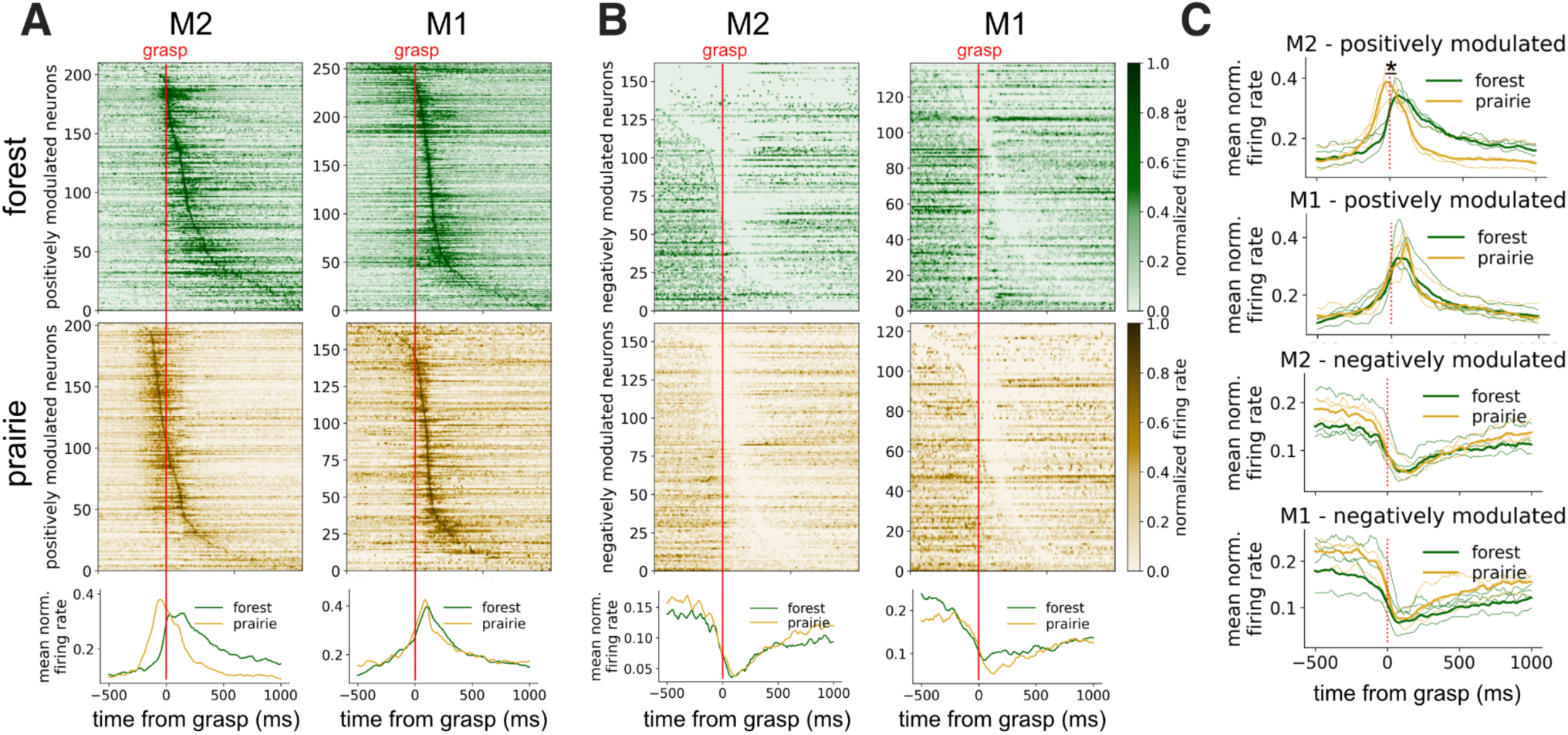
Neural activity in forest mouse M2 peaks closer to the grasp than prairie mouse. A. (*top*) Each heatmap shows normalized firing rates for all positively modulated units recorded in a single region in a single example animal, averaged across successful trials (forest mouse sample recorded over three sessions, prairie mouse animal recorded over five sessions). Data are aligned to the time of the grasp. (*bottom*) Average firing rate across all units for the example units shown. B. (*top*) Each heatmap shows normalized firing rates for all negatively modulated units recorded in a single region in a single example animal, averaged across successful trials(forest mouse sample recorded over three sessions, prairie mouse animal recorded over five sessions). Data are aligned to the time of the grasp. (*bottom*) Average firing rate across all units for the example units shown. C. Average firing rate across all positively or negatively modulated units for n=3 forest M2, n=4 forest M1, and n=3 prairie M1/M2. Thick lines represent the average of all units recorded in all animals, and thin lines represent the units from individual animals. Red lines mark the time of grasp. *p=0.049, two-sided Wilcoxon rank-sum test comparing the time at which the mean firing rate reached 90% of the maximum between subspecies (median [IQR]: NB, 537.0 [519.5–541.0], n = 3; BW, 432.0 [419.0–453.0], n=3; Z=1.96, r=0.80).

This difference in the timing of M2 neural activity could reflect differences in the reach-to-grasp behavior between subspecies. To test whether the difference in hand angle at grasp (Figure 4C-D) affects neural activity, we compared activity from the trials in which hand angles at grasp were most parallel to the platform (top quartile) to those where they were most perpendicular (bottom quartile) for each mouse. The most parallel prairie mouse hand angles overlapped with the distribution of forest mouse hand angles, and the most perpendicular forest mouse hand angles overlapped with the distribution of prairie mouse hand angles (see Figure S5F legend). We found no difference in the timing of the peak of activity in M2 between reaches with parallel and perpendicular hand angles within forest mice or prairie mice (Figure S5F), suggesting that the observed difference in activity is not driven by this behavioral difference. We next asked whether the subspecies difference we observe in M2 activity might be driven by the increased variability of forest mouse grasp position (Figure 4F-G) or overall reach trajectory (Figure S4I-J). We compared the most (top quartile) and least variable (bottom quartile) trials in each animal, as measured by distance from the median. We found that the least variable of forest mouse trials show variability that overlaps with the prairie mouse distribution, and the most variable prairie mouse trials overlap with the forest mouse distribution (see Figure S5G-H legends). We did not observe a difference in activity between the most and least variable trials for each mouse (Figure S5G-H), suggesting that the difference in neural activity is also not driven by this behavioral difference. We cannot rule out the possibility that the difference in neural activity is due to a difference in behavior that we cannot measure, such as fine digit movements or muscle activation missed by our kinematic analysis, nor can we rule out a difference in motor planning. However, our findings are consistent with the idea that circuit differences between forest and prairie mice result in differences in neural dynamics that enable more dexterous behavior in forest animals.

### Forest mice display more dexterous climbing behavior than prairie mice

In addition to its role in hand use, the CST has also been implicated in complex locomotion tasks, such as ladder walking^31,32^. Forest mice are thought to have evolved dexterous movement in response to the selective pressure to interact with a complex, three-dimensional environment, including by climbing. Indeed, a rich natural history literature reports better performance in climbing tasks in *P. maniculatus* subspecies that evolved in forests compared to their closest non-forest-dwelling relatives^39–42^. We therefore tested animals on a rod-crossing task, where they crossed a thin rod between two platforms, similar to movement across a thin twig, which has been documented in wild deer mice^61^, and similar to previous studies^40,41^. We made several adjustments to earlier versions of this assay to enable us to better measure animals’ climbing dexterity. We altered our assay to: (1) train animals to cross the rod to reduce the impact of the previously observed motivational difference between subspecies^40,55^; and (2) expose mice to progressively smaller and smaller rods to better identify the limits of the motor problems each subspecies could solve. We found that at the widest rod diameter, i.e., the easiest rod, both forest and prairie mice successfully cross the rod, although forest mice cross at a slightly higher velocity (Figure 6C). On narrower rods, we find that forest and prairie mouse performances diverge substantially (Figure 6A-F, Videos S7-18). Forest mice cross with higher velocities and spend most of their time moving and in an upright position, whereas prairie mice often stand still on the rod or hang upside down (Figure 6C-E). We also find that when animals flip upside down on any rod size, forest mice right themselves more quickly (Figure 6F). Finally, we observe that prairie mice more frequently fail to cross at all, with all (10/10) forest mice, but only 6 out of 11 prairie mice, succeeding in crossing the smallest rod without falling. Together, these data indicate that forest mice are more successful in the rod-crossing assay in a wider range of conditions and are thus more dexterous in both climbing and reaching.

**Figure 6:**
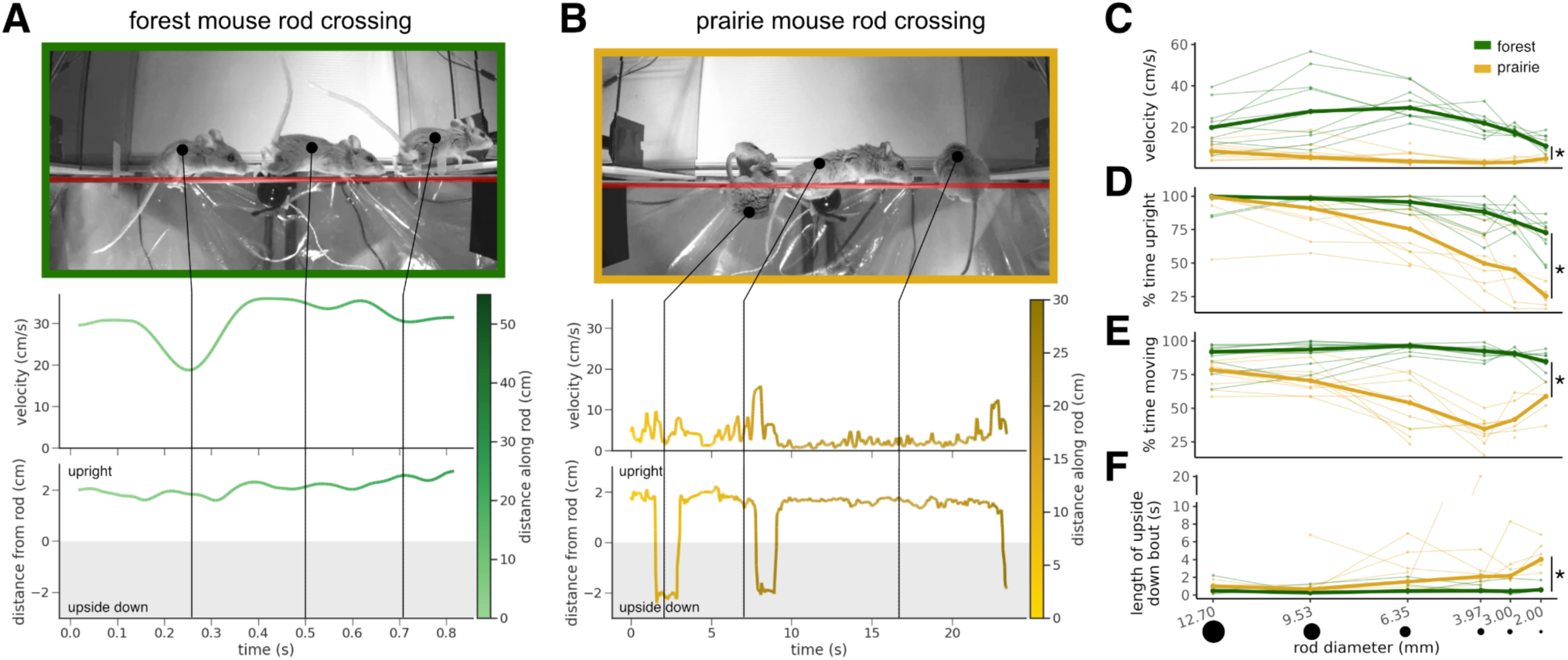
Forest mice show greater climbing dexterity than prairie mice. A/B. Example data from one forest (A) and one prairie (B) mouse crossing a 3.97mm rod. In this task, animals cross a rod between two platforms to receive a reward. Automated tracking of the animal’s back was used to determine the velocity and distance from the rod. The top image is made from video frames at the time points indicated by black lines. The rod is highlighted in red. C-F. Measures of rod-crossing performance across six different rod sizes. Animals crossed wider (easier) rods first and then were presented with progressively narrower rods. Thick lines show median performance, and thin lines show the performance of n=10 forest and n=11 prairie mice. C. Forest mice move across the rod faster than prairie mice (*p=9.6e-07, F=68.8, repeated measures ANOVA). D. Forest mice spend more time upright than prairie mice (*p=0.00019, F=25.1, repeated measures ANOVA). E. Forest mice spend more time moving than prairie mice (*p=3.6e-08, F=116.4, repeated measures ANOVA). F. Forest mice right themselves more quickly when they are upside down compared to prairie mice (*p= 0.0058, F=12.9, repeated measures ANOVA).

### Climbing dexterity is linked to corticospinal tract size

The evolutionary expansion of the corticospinal system, especially in the primate lineage, has been hypothesized to support dexterous movement, but testing this hypothesis has been difficult^19,20^. While CST or CSN manipulations support the role of the corticospinal system in dexterity^25–32^, how CST *size* or CSN *number* relates to dexterity remains unknown. To address this question, we took advantage of the interfertility of our forest and prairie deer mouse subspecies. We crossed forest and prairie deer mice and then crossed their progeny (F1s) to generate F2 hybrid mice. F2 mice have recombinant, shuffled genomes, with both forest and prairie elements (Figure 7A). This shuffling means that each individual F2 hybrid animal inherits different combinations of forest and prairie genes and, therefore, different neural phenotypes and behaviors. We hypothesized that if CST size is important for dexterous motor behavior, we should observe larger CSTs in more dexterous F2 animals. We assessed 51 F2 mice on the rod-crossing assay at three rod sizes. As expected, F2 animals showed a wide distribution of rod-crossing performance levels, as assessed by their velocity, time upright, time spent moving, and time to right themselves when they fall (Figure 7B-E). While we observe a correlation between F2 animals’ performances on the wider two rods (Figure S6A-B), this is not correlated with their performance on the narrowest rod (Figure S6C). This discrepancy suggests that crossing wider rods successfully requires different genetic, neural, and behavioral mechanisms than crossing the presumably more challenging, narrower rod. Therefore, we focused the rest of our analysis on performance on the narrowest rod, as we judged this to be most relevant to their dexterity and thus most likely to depend on CST size.

**Figure 7:**
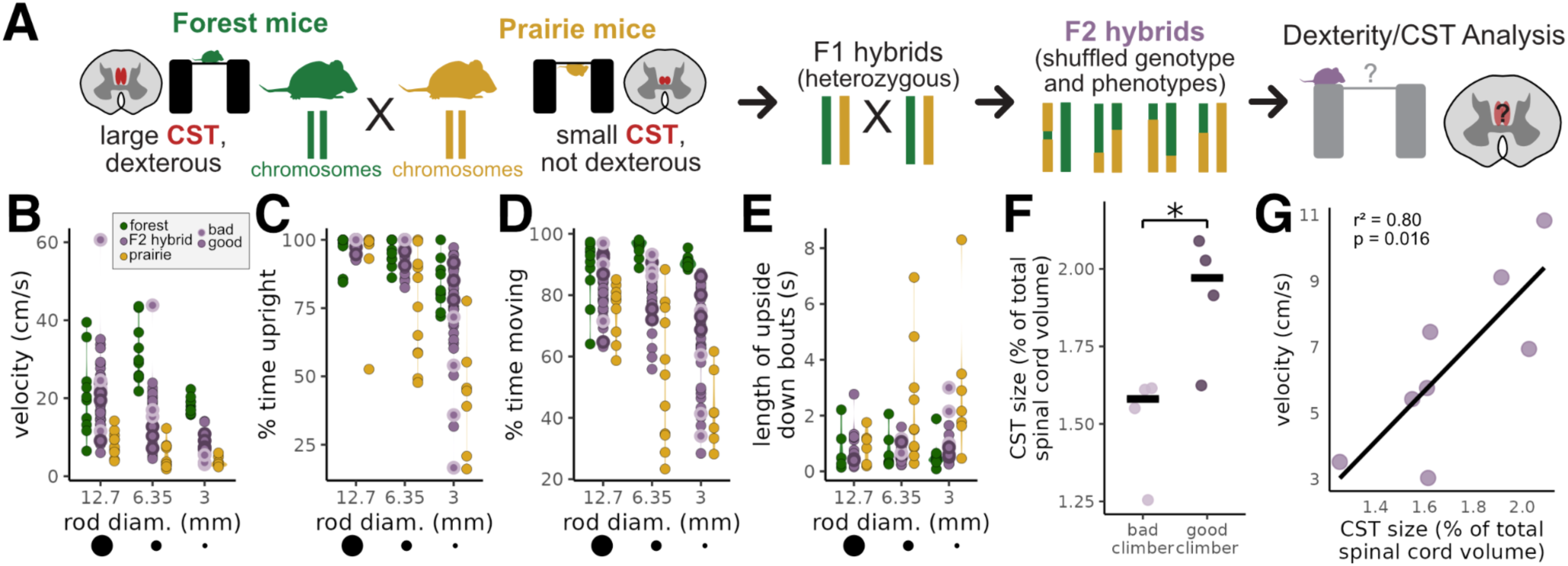
Corticospinal tract size is linked to climbing dexterity. A. Schematic of the F2 hybrid generation and testing. F2s were generated by crossing forest and prairie mice for two generations. F2 animals have shuffled genomes and phenotypes and can thus be used to separate subspecies’ phenotypes. B-E. Quantification of rod-crossing performance in F2 hybrid animals (forest/prairie data from Figure 6 shown for comparison). “Good” and “bad” climbers chosen for CST analysis are indicated. Data from n=41 F2 animals. F2s show a broad distribution of climbing dexterity. F. Good climber F2 animals have larger CSTs than bad climber F2 animals. Data are normalized by total spinal cord size. *p=0.029, two-sided Wilcoxon rank-sum test (median [IQR]: good climber, 0.020 [0.018–0.020], n=4; bad climber, 0.016 [0.015–0.016], n=4; Z=2.31, r=0.82). G. CST size is correlated with rod-crossing velocity in F2 animals. Pearson correlation: r^2^=0.81, t=3.3, df=6, p=0.016, n=8. Pearson correlation on residuals after accounting for morphological traits (weight, tail length, foot size): r^2^=0.81, t=3.4, df=6, p=0.014.

To determine whether CST size is linked to rod-crossing dexterity, we selected four “good” and four “bad” climbers based on their performance metrics. Interestingly, velocity and the fraction of time spent upright during crossing are not correlated (Figure S6D), indicating that these aspects of climbing dexterity are likely controlled by different neural and genetic mechanisms. Therefore, we identified “good climbers” as those in the top half of animals in both their velocity and time upright and “bad climbers” as those in the bottom half of animals in both measures. When we compared cervical CST volume between good and bad climbers, we found that good climbers have larger CSTs than bad climbers (Figure 7F). When we asked whether CST size correlates with rod-crossing velocity in these eight mice, we also found a correlation (Figure 7G), but we did not find a correlation between CST size and fraction of time spent upright (Figure S6E). While our data suggests that CST size impacts velocity but not time upright, suggesting different neural mechanisms impacting these two aspects of climbing, we do not have the power in this 8-animal CST-size dataset to completely rule out an association between CST size and time upright. Importantly, we find that tail length, weight, and hind foot size, traits commonly altered in arboreal animals and hypothesized to be involved in climbing performance^37,40,41,62^, are not correlated with velocity in a larger 41-animal dataset (Figure S6F-H). Furthermore, controlling for weight, tail length, and foot size, we still find a positive correlation between CST size and rod-crossing velocity. While we cannot exclude the possibility that another neural or morphological feature that confers climbing dexterity is genetically linked to CST size, the most parsimonious explanation for these data is that a larger CST, and likely more CSNs, supports climbing dexterity.

## Discussion

Across taxa—from birds to lizards to rodents to primates—arboreality has evolved repeatedly. Arboreal species or populations have convergently evolved morphological characteristics such as long tails, larger hands and feet, or longer whiskers, and behavioral factors such as differences in gait or hand use, often related to climbing dexterity^37,43,62,63^. However, less is known about neural adaptations to arboreality. We show that CSN number expansion may be a neural adaptation to life in a forested habitat. Consistent with this idea, primates, which also evolved arboreality, have expanded corticospinal systems, presumably to support increased dexterity^19,20,23^ (though see^17^). Indeed, dexterity may be an arboreal trait, though other environmental pressures likely also influence dexterity^64–66^.

In support of dexterity being an arboreal trait, we find that semi-arboreal forest mice are more dexterous in both a reach-to-grasp task and an ecologically relevant rod-crossing assay of climbing. The difference between forest and prairie mouse behavior is specifically in dexterity, that is, the ability “to adequately solve any emerging motor problem correctly, quickly, rationally, and resourcefully”, rather than in general motor coordination, that is, the ability to precisely control movements^33^. In a reach-to-grasp task, forest mice are more flexible in the conditions in which they can succeed, whereas prairie mice can learn to succeed only under specific circumstances and by using a stereotyped movement. Similarly, in a test of climbing performance, forest mice can dexterously climb narrow rods, whereas prairie mice can successfully climb only wider rods where quick responses to momentary loss of balance are less important. These two behaviors may be related; that is, manual dexterity may directly impact complex locomotion. On the other hand, neural mechanisms that enhance manual dexterity could also improve animals’ ability to solve motor problems beyond the hand. While we have not distinguished between these possibilities, we use F2 hybrid animals to demonstrate that CST size is linked to climbing dexterity. Thus, our data suggest that expansion of the corticospinal system likely confers dexterity in an ecologically relevant rod-crossing climbing task, supporting the idea that evolutionary expansion of the corticospinal system supports dexterous movement.

Variation in CSN number between forest and prairie mice could influence behavior through altering neural circuit dynamics. Consistent with this, we observe that forest animals with more CSNs also show a shift in the timing of cortical activity toward the time of the grasp during a reach-to-grasp task. Several lines of evidence suggest that this shift could be due to CSN number. First, we observe the shift in activity only in M2, where we observe more CSNs in forest compared to mice, but not in M1, where we do not. Second, a role for M2 CSNs in mediating specifically the dexterous grasping phase of the behavior is consistent with previous work^26,29^. Variation in CSN number could shift M2 neural activity towards the time of the grasp, as we observe, by altering the circuit to be more tuned to the sensory signal from the grasp. This could be due to a delay in activity caused by increased feed-forward inhibition during the reach due to reafference cancellation^67^. This reafference cancellation could be driven by CSN synapses onto inhibitory spinal cord interneurons^68^ and later overridden during the grasp by novel sensory input. Alternatively, it could be due to differences in local circuitry, for example amplification of sensory information via recurrent connections between CSNs and thalamus^69^. Finally, this shift could result from plasticity within the larger M2 sensorimotor circuit. For example, M2 inputs to the spinal cord could induce heterosynaptic plasticity, resulting in the strengthening of coincident ascending sensory feedback in response to increased motor signals^70–73^. In any case, a sensory-tuned circuit could enable enhanced flexibility in movement and increased dexterity. Given the potential role of M2 in planning^71^, we also note the possibility that the forest mouse strategy requires less planning, resulting in decreased early preparatory activity. Future work that specifically records from CSNs and manipulates them in forest and prairie mice will be necessary to fully understand how CSN expansion impacts neural dynamics.

The genetic variation driving the difference in CSN abundance likely affects embryonic and early postnatal developmental processes^74^. The neurons that become CSNs are born embryonically as a part of layer V. Then, a subset of these neurons is specified to become CSNs and directed to project to specific spinal targets via defined transcriptional programs^75–81^. Finally, some of these spinal-projecting neurons are pruned from the spinal cord postnatally and cease to be CSNs^82–85^. Differences in any of these steps between subspecies could result in the differences in CSN number that we observe, but given our results, it is likely that subspecies differences occur locally in M2 and S2, as we do not observe a broad increase in the number of layer V extra-telencephalic neurons or differences in CSN number in other cortical areas. An intriguing possibility is that the expansion of CSNs we observe is due to the expansion of a specific subpopulation of CSNs with a particular developmental origin^86,87^, synaptic partner in the spinal cord^68,73,88^ or general projection target in the brain or spinal cord^77–80,89^. The F2 hybrid deer mice provide a powerful system to investigate this question, given the ability to use forward genetics to potentially reveal the genetic basis of morphological and behavioral variation in future studies^90^.

The evolution of neuron number and connectivity, orchestrated through genomic changes, sets up the network structures that drive behavioral variation^91^. However, how altering neuron number impacts neural function to drive behavior is unclear. Evidence from artificial neural networks (ANNs) may provide some insight into this process. In many types of ANNs, increasing network size via increasing parameters (or sometimes neurons/synapses, more analogously) or increasing input data results in better performance, often described by scaling laws^92–98^. One potential explanation for this increase in performance is that increasing ANN size enables an increase in redundancy in the system, which results in faster learning as well as more robustness to noise^92,93^. This interpretation fits with some biological findings of robustness in behavior or circuit function^10,99^. Additionally, increasing ANN size, especially in large language models (LLMs), results in emergent properties of the network^94–98^. For example, with enough parameters, LLMs can overcome the rigidity caused by overfitting and regain flexibility while still fitting general trends, enabling them to implicitly learn language syntax^98^. While there are no direct biological analogies to this property, it is consistent with correlations between brain size and behavioral flexibility^100^, including our data. Here, we identify the deer mouse system as a promising model to answer a fundamental question in network function, relevant to both biological and artificial networks^91,101^: how does altering network structure, in particular neuron number, alter network function to achieve behavioral complexity?

---

**Supplementary Figure 1:**
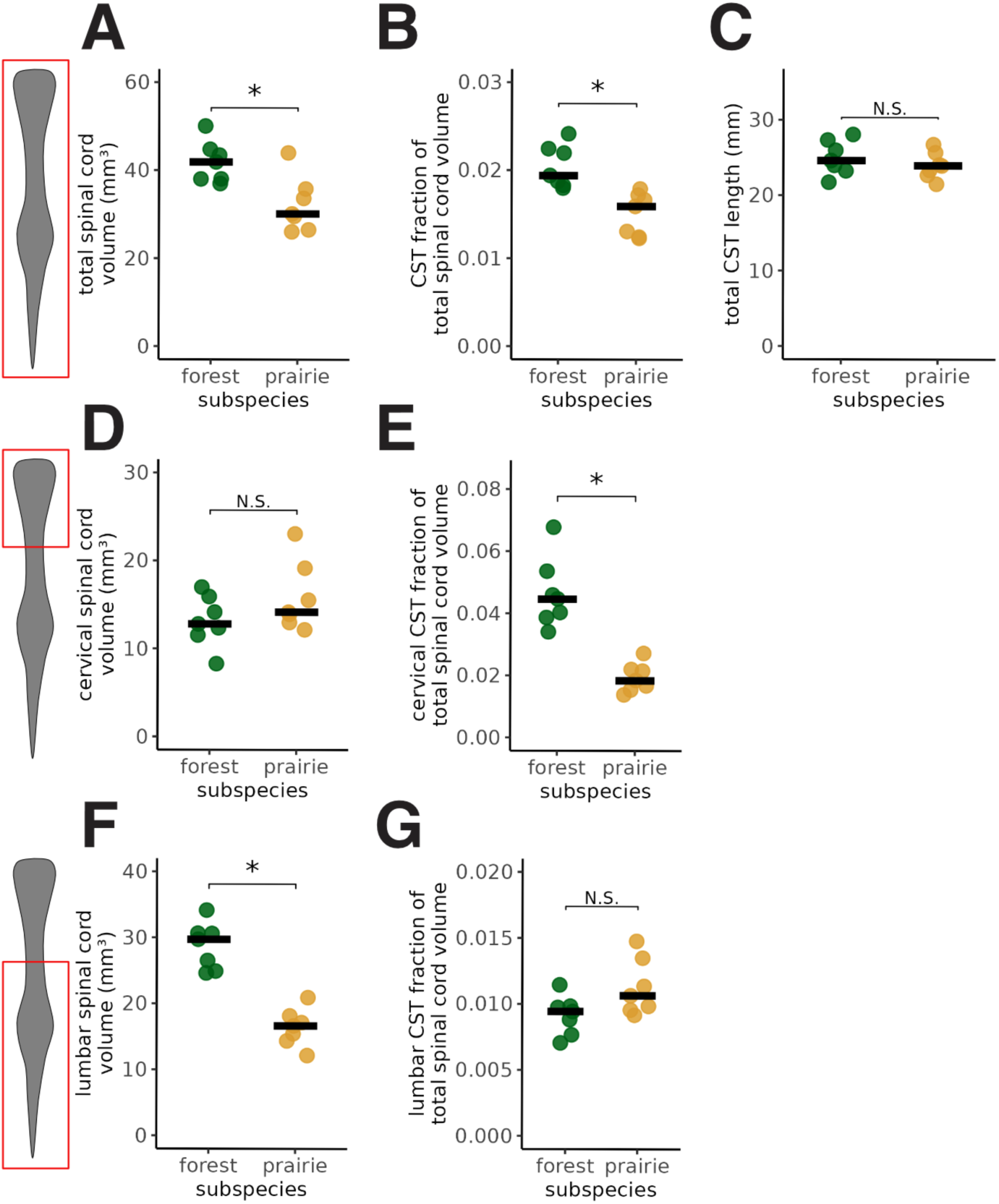
Corticospinal tract (CST) morphology. A. Forest mice have larger spinal cords. *p=0.011, two-sided Wilcoxon rank-sum test (median [IQR]: prairie, 30.117 [28.044–34.729], n=7; forest, 41.900 [38.028–44.111], n=7; Z=-2.49, r=0.67). B. The CST comprises a larger fraction of total spinal cord volume in forest mice. *p=0.001, two-sided Wilcoxon rank-sum test (median [IQR]: prairie, 0.016 [0.013–0.017], n=7; forest, 0.019 [0.019–0.022], n=7; Z=-3.13, r=0.84). C. Total CST length (from cervical to lumbar spinal cord) is not different between forest and prairie mice. N.S.: p=0.383, two-sided Wilcoxon rank-sum test (median [IQR]: prairie, 23.938 [23.007–24.877], n=7; forest, 24.615 [23.606–26.657], n=7; Z=-0.96, r=0.26) D. Cervical spinal cord volume is not different between forest and prairie mice. N.S.: p=0.32, two-sided Wilcoxon rank-sum test (median [IQR]: prairie, 14.151 [13.447–17.343], n=7; forest, 12.800 [11.944–15.033], n=7; Z=1.08, r=0.29). E. The CST comprises a larger fraction of cervical spinal cord volume in forest mice. *p=0.001, two-sided Wilcoxon rank-sum test (median [IQR]: prairie, 0.018 [0.016–0.022], n=7; forest, 0.045 [0.040–0.050], n=7; Z=-3.13, r=0.84). F. The lumbar spinal cord volume is larger in forest mice compared to prairie mice. *p=0.001, two-sided Wilcoxon rank-sum test (median [IQR]: prairie, 16.627 [14.891–17.639], n=7; forest, 29.725 [25.713–30.636], n=7; Z=-3.13, r=0.84). G. The CST comprises the same fraction of lumbar spinal cord volume in forest and prairie mice. N.S.: p=0.073, two-sided Wilcoxon rank-sum test (median [IQR]: prairie, 0.011 [0.010–0.012], n=7; forest, 0.009 [0.008–0.010], n=7; Z=1.85, r=0.50). All data are the same as in Figure 1 from n=7 animals of each subspecies.

**Supplementary Figure 2:**
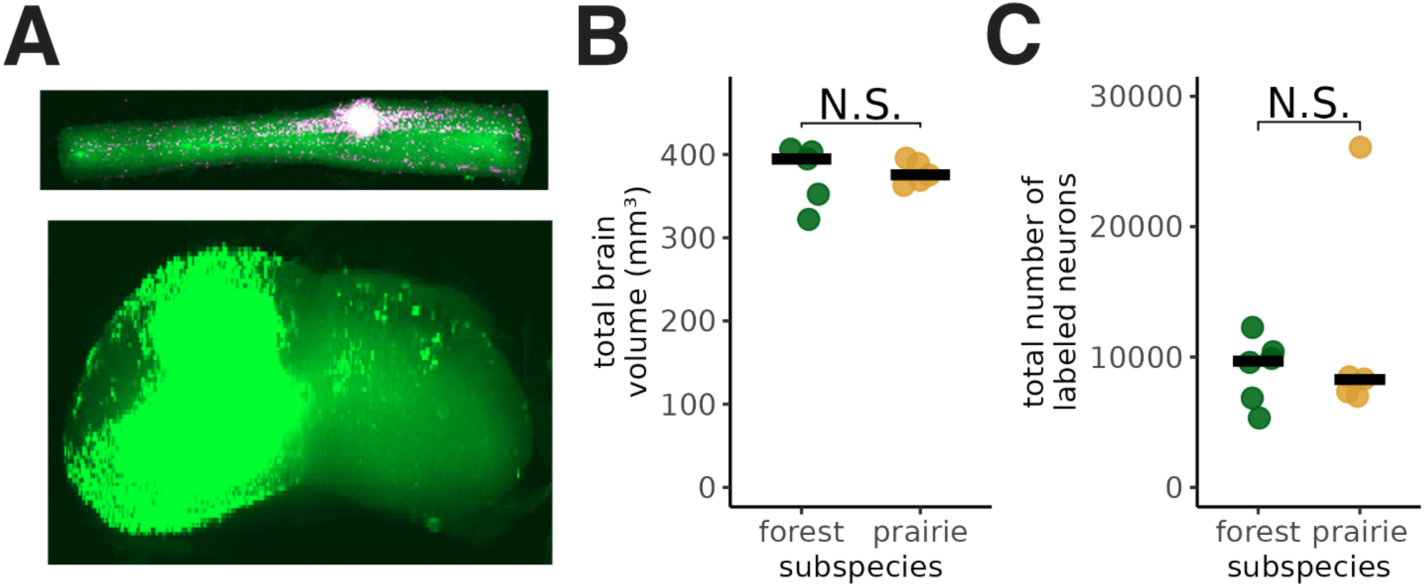
Retrograde viral tracing. A. Example light-sheet images from a spinal cord injection. *Top*: maximum intensity projection along the dorsal-ventral axis of cervical/thoracic spinal cord, showing the injection site. Magenta = anti-GFP staining, green = GFP signal. *Bottom:* maximum intensity projection along the anterior-posterior axis of the injection site from the same animal. Only GFP signal is shown. B. Forest and prairie mice do not differ in total brain volume, p=0.537, two-sided Wilcoxon rank-sum test (median [IQR]: prairie, 375.702 [369.664–389.593], n=5; forest, 399.094 [362.747–405.202], n=6; Z=-0.73, r=0.22). C. Forest and prairie mice did not differ in the total number of neurons labeled throughout the brain by retrograde viral injection, p=0.931, two-sided Wilcoxon rank-sum test (median [IQR]: prairie, 8275.000 [7321.000–8458.000], n=5; forest, 9682.500 [7474.000–10217.750], n=6; Z=-0.18, r=0.06).

**Supplementary Figure 3:**
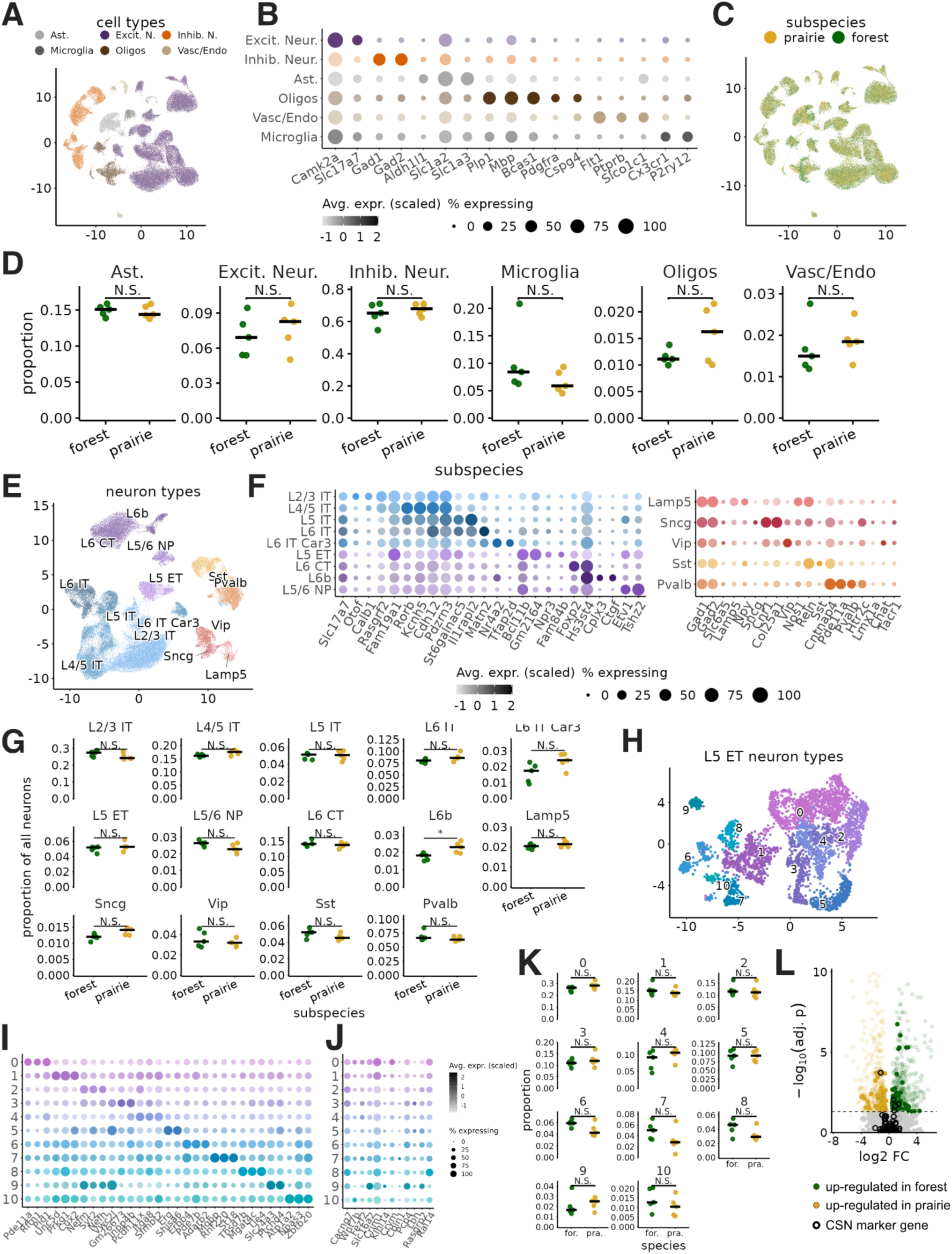
Single-nucleus RNA sequencing reveals no difference in L5ET or cortical neuron number between subspecies. A. UMAP plot of gene expression in all cortical nuclei isolated from forest and prairie mice, colored by their cell class, as determined by alignment to the BICNN database. Data are from 107,000 nuclei from n=5 animals from each subspecies. (Note that this is after integration – see Methods). B. Expression of established marker genes for cell class of the neural populations shown in A. C. UMAP plot of gene expression in cortical nuclei isolated from forest and prairie mice, colored by their subspecies. D. Percentage of nuclei per sample assigned to each cell class. N.S. = p>0.05, edgeR. E. UMAP plot of gene expression in neuronal nuclei (as defined in A) of both forest and prairie mice. Coloring is by their subclass. F. Expression of established marker genes for cell subclass of the neural populations shown in B. G. Percentage of nuclei per sample assigned to each cell subclass. * = p<0.05, N.S. = p>0.05, edgeR. H. UMAP plot of gene expression in L5ET nuclei (as defined in E) of both forest and prairie mice. Coloring is by unbiased clustering results. I. Expression of the top 3 marker genes for each cluster of L5ET neurons. J. Expression of known CSN marker genes from Golan et al, 2023, and Arlotta et al., 2005 for each cluster of L5ET neurons. K. Percentage of nuclei per sample in each cluster. N.S. = p>0.05, edgeR. L. Differential gene expression analysis of pseudo-bulk gene expression in L5ET neurons between forest and prairie mice. Dark circles mark CSN marker genes in Golan et al. Genes up-regulated in forest mice are enriched for CSN marker genes (p=0.0068, Fisher exact test), but those up-regulated in prairie mice are not (p=0.56, Fisher exact test). Genes differentially expressed between subspecies in all neurons are shown in a lighter alpha shade.

**Supplementary Figure 4:**
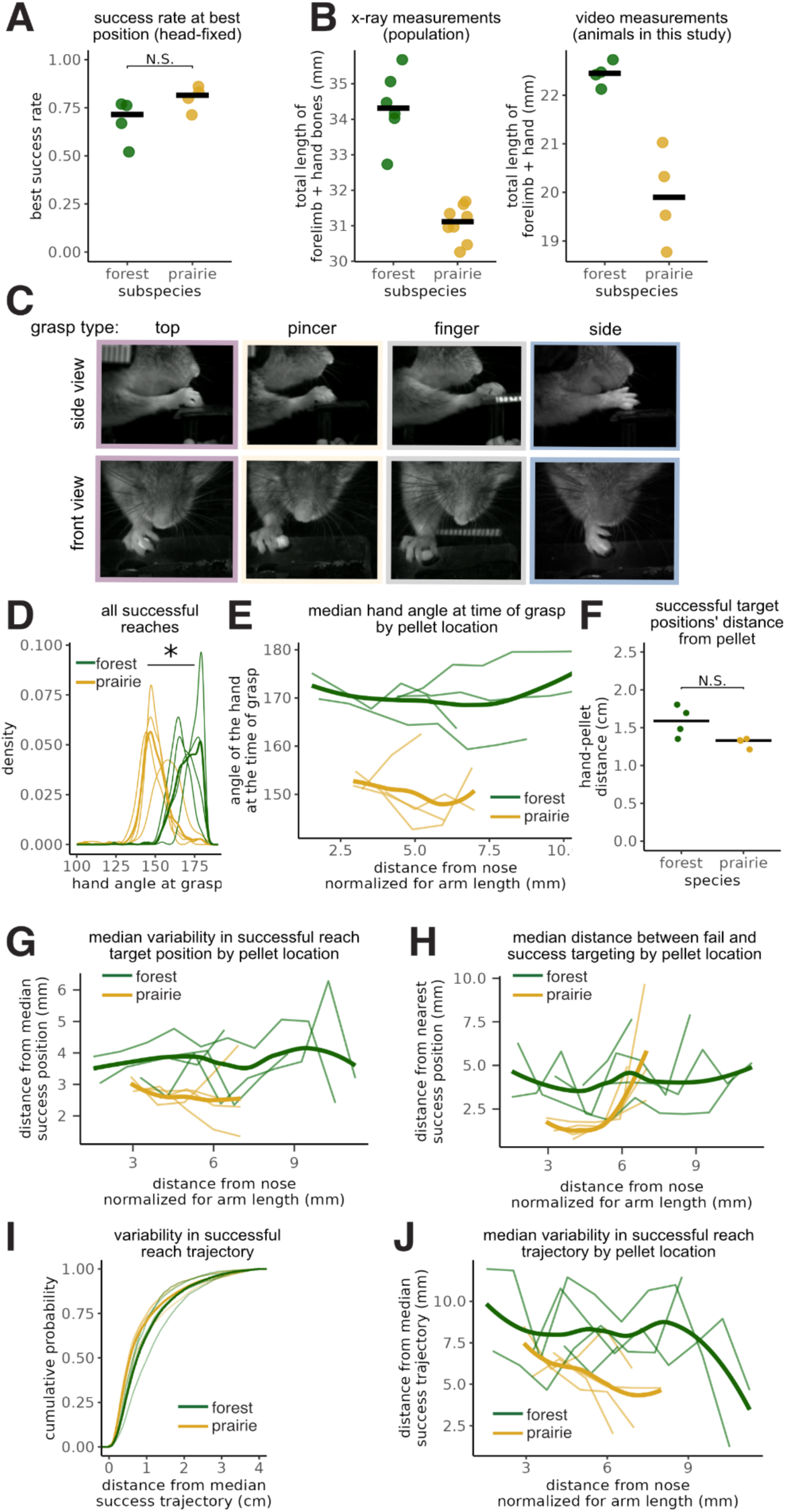
Manual dexterity in a reach-to-grasp task. A. In a head-fixed reach-to-grasp task (Figure 4B), forest and prairie mice achieve equivalent dexterity. The success rate at each animal’s most successful pellet position is not significantly different between subspecies (p=0.11, two-sided Wilcoxon rank-sum test (median [IQR]: prairie, 0.815 [0.778–0.837], n=4; forest, 0.715 [0.633–0.763], n=4; Z=1.58, r=0.61). B. Arm length measurements from x-ray data from Hager et al., 2020 (left) and those made from calibrated video images used in this study (right). We find similar differences between populations (x-ray: mean 3.2mm +/- 1.06 95% CI, video: mean 2.5mm +/- 1.47 95% CI), indicating that the values we used to normalize our data are reasonable. The difference in absolute lengths between x-ray and video measurements come from: (1) x-ray measurements sum bones that overlap and do not contribute to overall bone length, (2) x-ray measurements start at the shoulder socket, which is difficult to determine in video images. Video measurements were made blinded to the animal’s identity. C. Examples of the qualitative grasp types (Figure 4E). Images taken just as the animal begins to move the pellet. D. Forest mice use a more parallel, top-down grasp compared to their prairie counterparts. This is the same as Figure 4B, except using all successful reaches, not just those at the animals’ most successful positions. Lighter lines show distributions of individual animals (n=4 of each subspecies), darker lines show the distribution of the subspecies. Forest mouse hand positions are closer to parallel to the platform (top-down reach) at the time of grasp, *p=0.029, two-sided Wilcoxon rank-sum test on per-animal medians (median [IQR]: prairie, 148.5 [147.706–150.976]; forest, 169.89 [167.428–172.654]; Z=- 2.18, r=0.82). E. Forest mice use a top-down grasping strategy, and prairie mice use a side grasping strategy regardless of pellet position. Lighter lines show distributions of individual animals (n=4 of each subspecies), darker lines show the distribution of the subspecies. Forest mice showed higher hand angles; significant effect of subspecies after accounting for pellet position (linear mixed-effects model with mouse as a random intercept; χ²2=25.2, p=3.3e-06). F. Despite differences in hand morphology, forest mice do not show significant differences in the distance of their hand from the pellet during targeting (p=0.057, two-sided Wilcoxon rank-sum test (median [IQR]: prairie, 1.33 [1.298–1.339], n=4; forest, 1.589 [1.45–1.722], n=4; Z=-1.9, r=0.71; median distance between hand target position and pellet position plotted). G. Forest mice have greater variability in pellet targeting at all positions. This is the same data as in Figure 4G, divided by pellet position. Lighter lines show the median MAD (median absolute difference) for each pellet position; darker lines show the subspecies median. Forest mice showed higher MAD of grasp position; significant effect of subspecies after accounting for pellet position and hand-pellet distance, as well as their interactions (linear mixed-effects model with mouse as a random intercept; χ²₄=20.9, p=0.00033). H. Forest mice have greater variability in targeting the pellet even when accounting for pellet position. This is the same data as in Figure 4H, divided by pellet position. Lighter lines show the median distance between failure and successful reaches for each pellet position; darker lines show the subspecies median. Forest mice showed greater fail-success distance and subspecies differed in their dependence on pellet position (linear mixed-effects model with mouse as a random intercept; overall subspecies effect: likelihood ratio test, χ²₂=62.4, p=2.9e-14; subspecies × pellet distance interaction: β=-0.10, t=-7.05, p=2.1e-12). I. Cumulative distribution plot showing, for each successful reach attempt, the variability of 3D reach trajectory, i.e., the median absolute deviation or the distance of each trajectory from the median successful 3D trajectory. Light lines show individual animals, dark lines show subspecies distributions. Forest mice showed greater variability; significant effect of subspecies on trajectory and significant pellet position x subspecies interaction (linear mixed-effects model with mouse as a random intercept; χ²2=6.48, p=0.011; subspecies x pellet distance interaction: β=0.097, t=22.1, p<2.0e-16). J. Forest mice have greater variability in their trajectories regardless of pellet position. This is the same data as (I) divided by pellet position. Lighter lines show the median distance from the median trajectory for each pellet position; darker lines show the subspecies median. Same data and statistics as in (I).

**Supplementary Figure 5:**
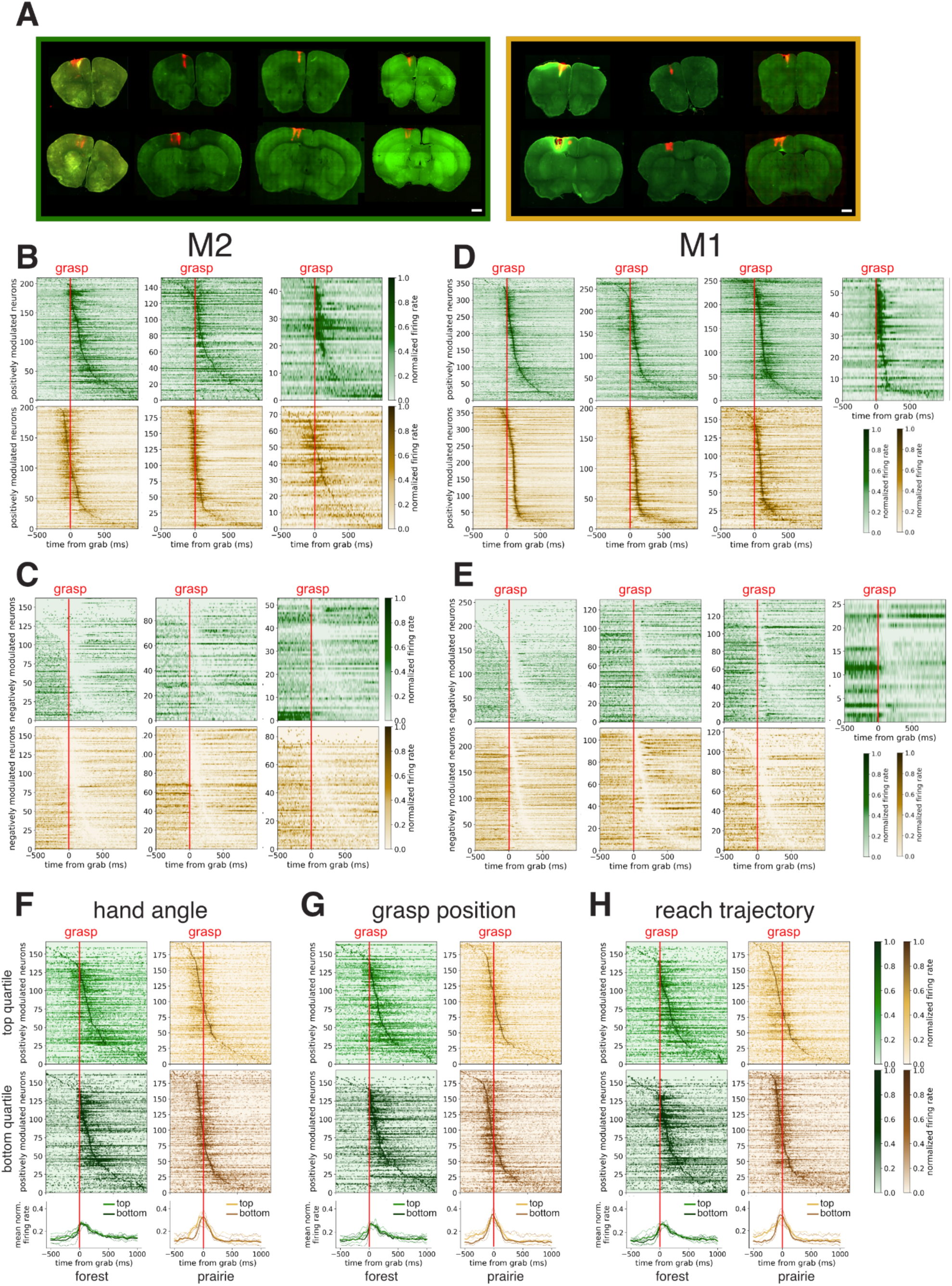
Neural recording during a reach-to-grasp task. A. Sections from forest (left) and prairie (right) deer mouse brains showing the location of the neural recording probe in each animal from DiI staining. Each column is one animal. The top row is M2, and the bottom row is M1. Scale bar is 1mm. Recordings from M2 in the third forest animal were discarded due poor signal-to-noise ratio. B-E. Each heatmap shows, for one animal, normalized firing rates for all positively (A/D) or negatively (C/E) modulated units recorded in a single region averaged across successful trials. Data are aligned to the time of the grasp. Data from M2 (B/C) or M1 (C/E). B/C. Data are from 3 sessions for each forest mouse and 5,4,5 sessions, respectively, for each of the prairie mice. D/E. Data are from 3,3,4,3 sessions for each of the forest mice, respectively, and 5,4,5 sessions, respectively, for each of the prairie mice. F. *(top)* Heatmaps of M2 neural activity for a representative animal of normalized firing rates for all positively modulated units averaged across the trials within the top 25% highest (i.e., most “top” grasp) hand angles or bottom 25% lowest (i.e., most “side” grasp) hand angles. *(bottom)* Average firing rate across all positively modulated units in M2 for n=3 forest and n=3 prairie mice across either the top or bottom quartiles, as indicated. Thick lines represent the average of all units recorded in all animals, and thin lines represent the units from individual animals. The mean hand angle (+/-SE) of top quartile trials was 169+/-0.7, 157+/-0.5, 172+/-0.7 for each of the prairie mice and 178+/-0.5, 173+/-1.2, 168+/-2.1, for each of the forest mice. The mean hand angle of bottom quartile trials was 154+/-0.4, 142+/-1.2, 157+/-0.8 for each of the prairie mice, and 169+/-1.4, 160+/-2.1, 149+/-0.8 for each of the forest mice. G. As in F, but for the top and bottom quartiles of variability of hand position at grasp, with the top quartile representing positions furthest from the median and the bottom representing positions closest to the median. The mean variability of top quartile trials was 0.45+/-0.1, 0.56+/-0.05, 0.36+/-0.1 for each of the prairie mice and 0.59+/-0.03, 0.74+/-0.03, 0.78+/-0.04, for each of the forest mice. The mean variability of bottom quartile trials was 0.12 +/-0.007, 0.10+/-008, 0.12+/-0.7 for each of the prairie mice, and 0.22+/-0.01, 0.17+/-0.01, 0.16+/-0.01 for each of the forest mice. H. As in F, but for the top and bottom quartiles of variability of 3D reach trajectory, with the top quartile representing positions furthest from the median and the bottom representing positions closest to the median. The mean variability of top quartile trials was 1.03+/-0.03, 1.86+/-0.10, 1.96+/-0.22 for each of the prairie mice and 1.96+/-0.16, 1.86+/-0.1, 2.15+/-0.26, for each of the forest mice. The mean variability of bottom quartile trials was 0.34+/-0.01, 0.52+/-0.4, 0.6+/-0.03 for each of the prairie mice, and 0.52+/-0.02, 0.78+/-0.9, 0.82+/-0.05 for each of the forest mice.

**Supplementary Figure 6:**
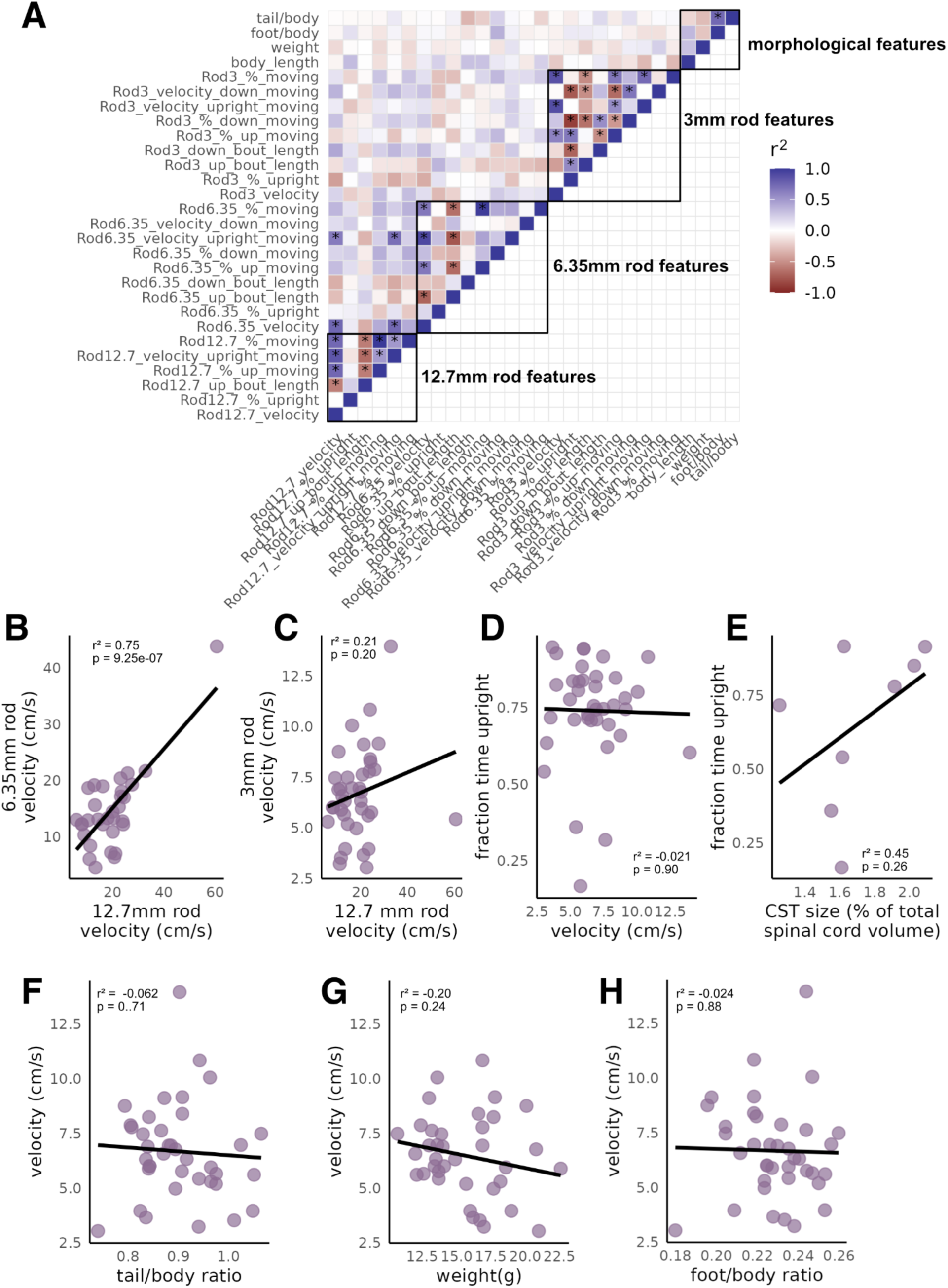
F2 hybrid mice. A. Correlation matrix of behavioral and morphological features in F2 animals. *=q <0.05, FDR-corrected p-values on r^2^ from a Perason correlation. Data are from n=41 F2 animals that had both morphological and behavioral measurements taken. B. Performance on the larger two rods is significantly correlated (data same as in A). Pearson correlation: r^2^=0.75, t=6.14, df=30, p=9.2e-07, n=32. C. Performance on the largest and smallest rods was not significantly correlated (data same as in A). Pearson correlation: r^2^=0.21, t=1.3067, df=36, p=0.20, n=38. D. On the smallest rod, velocity and the percentage of time spent upright are not significantly correlated (data same as in A). Pearson correlation: r^2^=-0.021, t=-0.12506, df=36, p=0.9, n=38. E. Fraction of time upright on the rod does not significantly correlate with CST size. Pearson correlation: r^2^=0.45, t=1.23, df=6, p=0.26. Data are from n=8 F2 animals in which we measured CST size. F. Velocity is not correlated with tail length (normalized to body size, data same as in A). Pearson correlation: r^2^=-0.062, t=-0.37508, df=36, p=0.71, n=38. G. Velocity is not correlated with weight (data same as in A). Pearson correlation: r^2^=-0.020, t=-1.20, df=36, p=0.24, n=38. H. Velocity is not correlated with hind foot length (normalized to body size, data same as in A). Pearson correlation: r^2^=-0.024, t=-0.15, df=36, p=0.88, n=38.

## Supplementary Videos

Videos S1-S6 are of animals performing the reach-to-grasp task. They are slowed down 10x. Each view is from a different synced camera.

Video S1: Example of a forest mouse grasping at the median forest mouse hand angle relative to the platform (171.6 degrees).

Video S2: Example of a forest mouse grasping at the median forest mouse hand angle relative to the platform (149.5 degrees).

Video S3: Example of a top grasp.

Video S4: Example of a side grasp.

Video S5: Example of a pincer grasp.

Video S6: Example of a finger grasp.

Videos S7-S18 are of animals performing the rod-crossing assay.

Video S7: Forest mouse crossing a 12.7mm diameter rod.

Video S8: Prairie mouse crossing a 12.7mm diameter rod.

Video S9: Forest mouse crossing a 9.35mm diameter rod.

Video S10: Prairie mouse crossing a 9.35mm diameter rod.

Video S11: Forest mouse crossing a 6.35mm diameter rod.

Video S12: Prairie mouse crossing a 6.35 mm diameter rod.

Video S13: Forest mouse crossing a 3.97mm diameter rod.

Video S14: Prairie mouse crossing a 3.97mm diameter rod.

Video S15: Forest mouse crossing a 3mm diameter rod.

Video S16: Prairie mouse crossing a 3mm diameter rod.

Video S17: Forest mouse crossing a 2mm diameter rod.

Video S18: Prairie mouse crossing a 2mm diameter rod.

## Methods

### Animals

We used two subspecies of deer mice: *Peromyscus maniculatus bairdii* and *P. m. nubiterrae*, as representatives of prairie and forest ecotypes, respectively. Forest mice were descendants of a colony founded by wild-caught deer mice trapped in Rector, PA, in 2010^37^. Prairie mice were descendants of a colony at the Peromyscus Genetic Stock Center (University of South Carolina), which are descended from wild-caught mice originally captured in Ann Arbor, MI, in 1948. Colonies were maintained using an outbred breeding scheme to maximize genetic diversity. Animals used in this study were of both sexes and between 3 months and 1 year of age. We used animals from different families (i.e., did not use only siblings/littermates) in all experiments to prevent confounding differences between subspecies with differences among families.

At Harvard University, mice were housed at 23°C on a 16:8 h light:dark cycle in standard mouse cages with corncob bedding, cotton nestlet, Enviro-Dri, and a red tube. Mice were fed lab mouse chow (LabDiet Prolab Isopro RMH 3000 5P75) ad libitum. At the University of North Carolina School of Medicine (UNC), mice were housed at 23°C on a 12:12h light:dark cycle in standard mouse cages with corncob bedding, cotton nestlet, and a red hut. Mice were fed lab mouse chow (LabDiet 5V5R - PicoLab® Select Rodent 50 IF/6F) ad libitum. All experiments were conducted in accordance with Harvard IACUC protocol 11-05 and UNC School of Medicine IACUC protocol 23-198.

### Spinal cord/brain clearing and antibody staining

For CST staining and retrograde tracing experiments in forest and prairie subspecies, animals were transcardially perfused with PBS with heparin (20U/mL) followed by 4% paraformaldehyde (PFA) in PBS. Brains were dissected out of the skull, and spinal cords were dissected using hydraulic extrusion^102^, and then post-fixed overnight in 4% PFA. Samples were cleared using a solvent-based method (adapted from previous work^103^). Briefly, they were dehydrated at 4°C using a methanol gradient with a minimum of 30 minutes at each step (20%, 40%, 60%, 80%, 100%) in B1n buffer (0.3 M glycine, 0.1% Triton X-100, 0.01% sodium azide, pH 7). They were then placed in dichloromethane overnight at 4°C and then rehydrated using the reverse methanol gradient. The spinal cords were then placed in B1n buffer overnight. Brains were further cleared using the uniclear method^104^ after the solvent-based method. They were incubated shaking at 37°C for 1 week in SiBP buffer (0.2mM, 0.08% SDS, 16% 2-methyl-2-butanol, 8% 2-propanol in water), changing the buffer after 3 hours, 6 hours, and then daily for the next week. They were then placed in B1n buffer overnight.

To stain, samples were incubated with antibody in PTxw buffer (PBS, 0.1% Triton X-100, 0.05% Tween20, 0.01% sodium azide) for 3-7 days (spinal cords) or 10-14 days, with a refresh in antibody after 5-7 days (brains), rotating at room temperature. They were then washed in PTxw and incubated with 1:200 secondary donkey-anti-mouse/rabbit-AlexaFluor647 antibody for 3 days (spinal cords) or 10 days (brains). Antibodies used were: A-7 mouse-anti-PKCy (sc-166451) at a concentration of 1:200, rabbit-anti-GFP (Invitrogen, A-11122) at 1:1000. They were then washed again with PTxw.

To image, spinal cords were refractive index matched in RI solution (a 60:40 mix of OptiPrep and 2,2-thiodiethanol with 0.6g/mL Histodenz) overnight. To fit into the microscope, spinal cords were cut in half at the thoracic region before imaging. Each half was mounted in a glass capillary tube. They were then imaged in RI solution in a Zeiss Lightsheet.7 microscope with a 5x objective, and a 1.525 refractive index. Brains were refractive index-matched and imaged in EasyIndex from LifeCanvasTechnologies (EI-500-1.52), also in a Zeiss Lightsheet.7 microscope with a 5x objective, and a 1.52 refractive index.

For F2 experiments, we modified our clearing protocol due to changes in university regulations on the use of dichloromethane. We cleared samples using the Life Canvas Technology passive clearing kit (C-PCK-500-1.52) following the instructions of the manufacturer. We note that our previous method altered spinal cord volume more than this method, and absolute volumes are not comparable between methods. Antibody staining was performed as described above. Spinal cords were imaged in EasyIndex on a Zeiss Lightsheet.7 microscope with a 5x objective and a 1.52 refractive index. All lightsheet images were acquired using ZenBlack software.

### CST volume quantification

To quantify the corticospinal tract (CST) size, images from the microscope were first stitched together in Arivis software. Then, using Arivis, we developed a pipeline that identifies the CST using a machine learning model. We then divided the CST into segments along the anterior-posterior axis of the spinal cord and quantified volume in those segments. Total spinal cord volume was quantified by thresholding the green autofluorescence signal. To compare between animals, the distance along the spinal cord was scaled using the length of the thoracic segment (we did not use total length due to the potential for differences in where the cut between the brain and spinal cord was made during dissection). For plotting, data were aligned such that T1 is the 0 point in all scaled data. Data were smoothed using the loess function in R with f set to 0.5. Statistical testing for volume differences was performed in R using a linear model: CST volume ∼ subspecies + age + total spinal cord volume + sex. We observed significant differences only for subspecies, but not other factors. Rank-sum tests were performed using the wilcox.text function in R.

### Dorsal funiculus axon immunostaining, confocal imaging, and quantification

Animals were perfused and spinal cords fixed as described above. Spinal cords were then mounted in agarose and sectioned on a vibratome at 75 μm thickness. We note that we found it is imperative not to freeze the sample to enable observation of axons. After sectioning, spinal cords were blocked in normal donkey serum, immunostained as free-floating sections with 1:200 A-7 PKCy antibody, with gentle shaking at room temperature overnight. They were then washed with PBS and stained with AlexaFluor 555 or 647 secondary antibody and Green FluoroMyelin (F34651) at 1:300 for 3 hours at room temperature. The dorsal funiculus of each section was imaged on a Zeiss LSM-900 confocal microscope with a Plan-Apochromat 40x/1.3 oil DIC (UV) VIS-IR objective with the following settings: 488 laser at 0.10% and 0.88 AU pinhole, 640 laser at 0.6% and pinhole of 0.78 AU or 561 laser at 0.2% power with 0.87 AU pinhole, scan speed of 4, gain of 650V, and averaging of 2. Images were acquired using ZenBlue software.

Image tiles were stitched in ZenBlue software and then exported as TIFF files. The CST area was manually segmented out using thresholding of the PKCy signal, with the experimenter blinded to the subspecies of the animal. Then, using custom Python code, 10-20 random regions of interest (ROIs), either 500x500 pixels or 250x250 pixels within the CST were selected for axon counting. Due to injury to the tissue during sectioning and staining, not all ROIs had quantifiable axon signal, and ROIs without such signal were manually filtered out with the experimenter blinded to the subspecies of the animal, ensuring that at least 5 500x500 ROIs or 10 250x250 ROIs per animal remained. The FluoroMyelin signal on good ROIs was inverted, and 6 ROIs from across samples were used to train a custom Cellpose^105^ model. The Cellpose model was then run on all good ROIs and manually checked to confirm good axon detection. We then filtered out any putative axons detected by Cellpose smaller than 0.1um^2^ as likely noise. We chose this value as the division between two peaks in a histogram of putative axon size. Total axon number for each animal was calculated by averaging the number of axons per micron in all of the ROIs counted, and then multiplying by the area of the CST. Statistical analysis was performed in python using the scipy.stats package.

### Electron microscopy preparation, imaging, and quantification

Animals were transcardially perfused with cold Ames buffer followed by cold 2% paraformaldehyde/2.5% glutaraldehyde in 0.1M cacodylate pH 7.4 with 0.04% calcium chloride. Spinal cords were dissected out and then incubated overnight in the same solution. After washing, spinal cords were coronally sectioned on a vibratome, followed by fixing of the sections in 3% glutaraldehyde in 0.1M cacodylate pH 7.4 with 0.04% calcium chloride. The dorsal funiculus was then dissected out and used in the following steps. After washing in water and maleate buffer pH 5.2, sections were incubated up to 24h in 1% uranyl acetate and washed again in a maleate buffer, followed by water. Next, they were incubated in lead aspartate for 3h at 60°C, washed with water, and dehydrated in an ethanol gradient at 4°C. They were then incubated in 100% ethanol at room temperature, followed by propylene oxide, and then embedded in LX112 resin and sectioned on a microtome to create ultrathin sections. Immediately before imaging, they were treated with 0.2% lead citrate. Sections were imaged on a JEOL 1200EX transmission electron microscope at 4800x. We imaged the ventral-most region of the dorsal funiculus and imaged at least 12 regions per sample.

To quantify the images, we used AxonDeepSeg^106^ to segment myelin and axons, following instructions for the published version with an additional myelin cutter model (https://github.com/axondeepseg/nn-axondeepseg) to better separate individual axons. We manually inspected and adjusted a subset of the images (10%) drawn from all animals. We compared the results from our manually refined dataset to the whole dataset and observed no differences, so we did not perform the manual refinement for the final data analysis (which enabled us to quantify all 12 images per animal). We used AxonDeepSeg to generate tables of axon size, myelin thickness, and g-ratio that we then analyzed using custom R scripts.

### Retrograde labeling surgeries

Animals were anesthetized with isoflurane for the duration of the surgery. After shaving the back, an incision was made in the skin. After tearing the muscle with forceps, the spinal column was clamped and stabilized using the Narishige spinal cord clamp (STS-A). The T1 vertebrae was located, and cervical injections were made 2-3 vertebral segments anterior to T1. The dura was removed from the spinal cord using a needle. Using a stereotaxic manipulator holding a pulled glass pipette, 25nL of virus was injected 1mm lateral from the midline (as defined by the central artery) at three depths: 500nm, 300nm, 200nm, for a total of 75nL injected. Each injection was done over 3 minutes with 3 minutes following the injection without moving the needle. Gelfoam was added to the spinal column to aid in healing, and the wound was closed with wound clips. Animals were perfused (see above) 3 weeks following the surgery. Animals were injected with SL1-AAV^49^ with H2B-linked green fluorescent protein (GFP) under the CAG promoter. We imaged the injection site in cleared spinal cords to confirm that it was in cervical segments 5-8. We eliminated 2 samples where injections were in thoracic spinal cord.

### Brain-wide retrograde labeling counting

Image tiles were stitched in Arivis software and then exported as TIFF files. TIFF files were stitched together into full 3D brain images and downsampled 3x using ANTS^107^. Images were then subdivided into smaller, overlapping 3D cubes (512x512x512 voxels) for counting. An ilastik^108^ model using a subset of the data taken from all brains was trained on the far red channel to identify GFP+ nuclei, background, and noise signal. We eliminated two samples at this point (one of each subspecies) due to poor perfusion, resulting in pervasive signal from blood vessels that was indistinguishable from neurons by our ilastik model. Using custom Python scripts, the GFP+ nuclei masks returned from ilastik were stitched together and assigned labels (where each nucleus has a unique label). To compare the same brain regions between animals, the green channel from the brain imaging (mostly autofluorescence because the clearing process quenches most of the GFP signal) was registered to a template brain generated by averaging the green channel autofluorescence signal of 40 non-injected brains imaged with a lightsheet microscope. Label files were then further downsampled, again using custom Python scripts, preserving the total count, and this downsampled image was transformed into template brain space using the registration from the green channel signal. Cortical regions were annotated based on manual adjustment of the Allen brain annotations registered to our template brain. To normalize for viral injection variation, we normalized nuclei counts by the total number of labeled nuclei in the brain. To ensure that this baseline count was not excessively affected by noise, we masked ventricles and other locations where there was noise common across samples. Furthermore, we created an average of the signal across samples (which was made up of samples that had injections on both sides of the spinal cord), and we masked areas where signal was either non-symmetrical or did not pass a threshold. We then used these regions to quantify the total number of neurons in each brain. We eliminated S2 from the analysis of one animal because of tissue damage in this region, resulting in no identifiable GFP signal. Downstream data analysis and plotting were performed using custom Python and R scripts.

### Single nucleus RNA-sequencing

Five animals of each subspecies were sacrificed using carbon dioxide, followed by decapitation. Brains were immediately removed, and the cortex was dissected from the rest of the brain. The anterior half of the cortex was extracted from both hemispheres for each animal (this includes all of sensorimotor cortex, as well as other areas) and flash-frozen in liquid nitrogen. Samples were stored at -70°C until further processing. Nuclei were extracted as described^109^ except that the high sucrose buffer had 1.2M sucrose. Sequencing libraries were prepared using the Parse Evercode WT Mega kit (along with other samples for different projects) according to the instructions of the manufacturer.

Samples were sequenced on a NovaSeq S4. Data were demultiplexed using python-based pipelines from Parse and aligned to the *P. maniculatus bairdii* genome^110^. To avoid confounds from mapping to the genome of one of the subspecies, before alignment, genomes were “personalized” to each subspecies, using a method similar to the generation of the *P. polionotus* genome^111^. Briefly, we aligned our data to the *P. maniculatus bairdii* genome using STAR aligner^112^. Then we performed variant calling for the individuals we sequenced for each subspecies using GATK. We then incorporated variants back into the final genomes to which we mapped reads. We did the final alignment using STAR aligner as a part of the Parse pipeline. Cells that passed a UMI threshold determined automatically by the Parse preprocessing script were carried on to the next steps.

Data were analyzed as a cell x gene count matrix in Seurat version 4.3.0^113^. Data were normalized with the SCTransform method and integrated across samples and subspecies using harmony^114^. Data were then clustered using the default Seurat settings with the FindClusters and FindNeighbors functions. Data were then mapped to the BICNN mouse motor cortex dataset^50^ using Azimuth to identify cell types. Clusters with low prediction scores for the mapping (median prediction score < 0.75 out of 1.0) were further assessed by finding marker genes. If clusters showed markers of multiple larger cell types (e.g., neurons and oligodendrocytes) or if they appeared to be non-cortical cell types (e.g., from dissection error), they were removed from the dataset. After removing this contaminating signal, this alignment resulted in 96% of cells with a prediction score greater than 0.75 when mapping to gene classes. The remaining clusters were then re-normalized, re-integrated, and re-clustered. We assessed differential abundance between subspecies for each cell type using edgeR^115^, as described previously^11^. We then subset neurons, re-normalized, re-integrated, and re-clustered, and used a combination of the Azimuth mapping and manual marker assessment to assign clusters to the BICNN cell types. At this stage, we eliminated two small clusters that had both strong excitatory neuron and strong oligodendrocyte markers. We again assessed differential abundance using edgeR. For alignment to subclasses, we found that 78% of neurons had an alignment prediction score greater than 0.75. When we took the subset of only L5ET neurons, we again renormalized, reintegrated, and reclustered (resolution = 0.3), and we performed differential abundance analysis as before. For pseudo-bulk differential gene expression analysis, we combined RNAseq reads for all L5ET neurons (or all neurons) by subspecies and sample. We then used DESeq2^116^ with default settings to calculate differential expression (DE). We eliminated genes that were DE in all neurons to focus specifically on genes DE in L5ET neurons. We compared those with marker genes for CSNs from the list of L5vsCSN genes described previously^52^.

### Reach-to-grasp behavior experiments and analysis

#### Freely moving reach-to-grasp experiments

Freely moving pellet reaching experiments were adapted from the protocol described previously^117^. Briefly, food-restricted animals were placed in a clear acrylic box with a slit and presented with a pellet in a small divot on a rotating platform 1.25cm from the front of the box. In the shaping phase, the slit was in the middle of the box. The shaping phase lasted until animals reached for 30 pellets 70% with a single hand. Animals were then moved to the training phase, in which they were placed into a box with the slit on the side of their preferred hand from the shaping phase. They were then presented with 30 pellets in each session and scored on how many of the pellets they successfully grabbed and ate. Animals were trained for 6 days. Animals were videotaped with FLIR Blackfly3 cameras at 500fps. Data was scored manually. Analysis and statistics were done using custom R scripts.

#### Head-fixed reach-to-grasp experiments

Head-fixed pellet reaching experiments were adapted from previous established protocols in *Mus musculus*^57,59,118^. Mice of either sex aged 16–50 weeks were fitted with head posts as described previously^59^, acclimated to head restraint, and water restricted (0.04-0.05 ml/g body weight) to 75-80% of their original body weight to encourage task participation. In each session, mice first learned to retrieve food pellets resting on a rotating table^57,59^, followed by a moveable vertical pellet dispenser that presents one pellet per trial for a limited period of time (3-5s; Sauerbrei et al., 2021). Each trial is video recorded using two high-speed cameras (FLIR, Flea 3, 500 fps) with manual iris and focus lenses, (Tamron 13VM1040ASIR 10–40 mm, f/1.4) one placed in front and one to the right side of the animal, illuminated with custom IR LED light sources, and calibrated for 3D tracking using DeepLabCut version 3.0.0^119,120^ with default settings. We tracked forelimb and digit movements first using a 2D DeepLabCut model trained and run using default settings, and then filtered using the ARIMA filter with ARdegree and MAdegree set to 1 and p_bound set to 0.2.

Triangulation was then run on the filtered data using default settings to create the 3D output. The cameras, acoustic cue, and pellet dispenser were controlled using Wavesurfer software (Adam Taylor, Janelia Scientific Computing; http://wavesurfer.janelia.org/) and a custom Arduino controller (Peter Polidoro, Janelia Experimental Technology).

#### Arm measurements

Arm measurements were taken by an experimenter blinded to the animal subspecies from video frames where the arm and fingers were fully extended. Lines were drawn from just before the shoulder (judging where the video showed movement), roughly below the posterior edge of the eye, to the tip of the longest finger, tracing the length of the arm. This was used to find the length of the arm in pixels. Pixels were converted into mm using a ruler roughly in the same plane of the frame as the arm (attached to the pellet platform). A similar procedure was used to measure the distance between the animals’ noses and the pellet position. This information was used to calculate the adjusted arm length, which was (distance from nose) - (arm length - minimum arm length of all animals), which essentially treats the distance for animals with longer arms as shorter by a value proportional to the length of their arms. For x-ray measurements, we used the length of the humerus + ulna + metacarpals + proximal and distal phalanges to get the total hand and arm length. We note that in our video data, we did not measure from the shoulder socket, as we did in x-ray data, and that x-ray data also is an overestimation of arm length due to overlap between the individual bones.

#### 3D video analysis

Success or failure was called manually. Similarly, grasp time was determined as the time when the animal first made contact with the pellet, as notated by manual observation. Hand angle was calculated from the DeepLabCut tracking data, taking a line between digit two and digit four and computing the angle between that line and the pellet platform. To compare similar parts of the grasp between animals and reduce noise in the angle measurement, the hand angle used in analysis was the angle at the time within 50ms before the grasp, where digits 2 and 4 were most spread apart (although we observed similar differences when using the grasp time exactly). To determine the frequency of each qualitative grasp type (top, side, finger, pincer), an observer blinded to the identity of the mouse classified each grasp (all grasps were shown in a scrambled order). To calculate hand position at the time of grasps, the 3D position from DeepLabCut tracking was used. The median success position used to calculate the median absolute difference (MAD) was the median x, median y, and median z from all successes for each animal for each behavioral session. Sessions with fewer than five successful reaches were discarded. To calculate pellet position, we tracked the pellet in DeepLabCut and took the median 3D pellet position before the animal reached for each day. To calculate the lift time (i.e., the start of the behavioral sequence), we first determined the times when the position of the wrist peaked in the y plane (up-down) and then defined the lift as the time before the peak where the acceleration started to increase. To calculate the fail position, we considered all positions between the lift time and 250ms after the lift and calculated the position at which the hand was closest to a position at which the animal successfully grasped the pellet in other trials. We again analyzed each behavioral session separately and eliminated sessions with fewer than five successful reaches. We further eliminated any reaches in which the hand was not within 4.2 cm from the pellet position, observing that those were mostly instances where the animal did not fully attempt to reach, as indicated by not fully raising the arm to the table. We determine this threshold value by taking the 99th percentile of the hand-pellet distances from successful reaches. The reach trajectory was calculated by taking the time between lift and grasp for each successful reach, interpolating the 3D location values so that each reach was on the same timeline, starting with lift and ending with reach. Then the Euclidean distance between each trajectory and the median trajectory for each behavioral session was calculated.

#### Statistical analysis

Data analysis was done using custom R and Python scripts. Rank-sum tests were performed using default settings of R’s wilcox.test() function. Linear regression was performed using the lmerTest package, fitting a linear mixed effects model with random intercepts for each mouse, with the following tests:

1. Hand angle: angle ∼ subspecies*pellet_position + (1|mouse)
2. Success position MAD: succ_position ∼ subspecies*pellet_position*pellet_hand_distance + (1|mouse)
3. Fail-success distance: fail_succ_dist ∼ subspecies*pellet_position + (1|mouse)
4. Distance from median trajectory: traj_dist ∼ subspecies*pellet_position + (1|mouse)

We assessed the overall effect of subspecies on each model by comparing models with and without subspecies terms using a likelihood ratio test.

### Neural recording experiments and analysis

At least one day prior to recording, two craniotomies were made over M1 and M2 within the following coordinates: M1, A/P: 0.0-0.5 mm, M/L: 1.0-2.0 mm; M2, AP: 1.0-2.0 mm, M/L: 0.8-1.3 mm. Depth targeting layer V: ∼0.4-1.2 mm from surface. Coordinates for recordings were determined based on the center of the CSN population in the retrograde tracing experiments. Electrophysiological recordings were performed similarly to prior work in *Mus musculus*^57,118^, using one 4-shank 384-channel Neuropixel 2.0 probe^56^ (IMEC, https://www.neuropixels.org/) per area (M1/M2). Data were spike-sorted using kilosort4^121^ and manually refined using the phy interface. Multi-units were removed from further analysis. Recordings were synced to the cameras, enabling us to match tracked behavior (see above) with neural activity. Neural data were binned (1ms) and smoothed using a 20-ms Gaussian kernel. Data were then normalized at the level of the unit using the following formula: (activity – minimum activity)/(maximum activity – minimum activity +2). This normalization was applied to ensure that each neuron contributed on a comparable scale, independent of absolute firing rate. We adopted min–max normalization for three primary reasons. First, it standardized activity distributions across sessions. Second, it allowed fair comparison across subspecies by placing all neurons on the same bounded scale. Third, given our relatively low sample sizes, min–max scaling stabilized variance estimates, reduced the influence of outlier neurons, and ensured that population-level analyses reflected consistent task-related dynamics. We isolated modulated neurons as those that had a significant (FDR-adjusted q < 0.05) difference in spike events in a 500 ms window centered 750 ms before the beginning of behavior, compared to a 500 ms window centered 250 ms after the beginning of the behavior. We further analyzed only modulated neurons. We identified the start of behavior as the lift time, determined from behavior tracking data. We first determined the times when the position of the wrist peaked in the y plane (up-down) and then defined ‘lift’ as the time before the peak where acceleration started to increase.

### Climbing behavior experiments and analysis

Animals were water-restricted to motivate participation in the task. Water was removed from home cages 1 day before the task, and animals were given at least 40μL of water per gram of body weight per day (mostly during the task, but they were supplemented if they did not drink enough). They were placed in an arena with two platforms separated by a 30cm rod with a plastic net underneath to catch them if they fell. Rods were stainless steel rods from McMaster Carr of the following diameters: 0.5 inch (½”, 12.7mm), 0.375 inch (⅜”, 9.53mm), 0.25 inch (¼”, 6.35mm), 0.15625 inch (5/32”, 3.97mm), 3mm, 2mm. Animals were trained over a ∼3-day period. First, they were trained to use the water-dispensing nose-poke on an enclosed platform. Next, they were exposed to the opportunity to cross to the other platform (forest and prairie mice) or the thickest rod (F2s) and were able to get water at either platform. Finally, they were exposed to a scenario, programmed using Bonsai^122^, in which they only received water when they successfully moved from one platform to another: water dispensers only turned on when the animal crossed an infrared beam at the entrance to each platform consecutively. Animals were given 1 day on each rod size. They failed if they fell off the rod more than 3 times in a 15-minute period or if they refused to cross the rod in that period. Animals who failed were given one more day to attempt that rod, but if they failed again, they were not given the next rod. We attempted to stop animals from crossing after 18 crosses per day (when they should have received their target water allocation for the day), but some animals performed slightly more crosses due to difficulty in removing them from the behavioral assay. Animals were removed from the behavior setup after 15 minutes, regardless of the number of crosses they achieved. Animals were videotaped using three synced Blackfly3 cameras recording at 120 fps: one from the top and two from the sides, angled up from the bottom. Video collection was done using campy (https://github.com/ksseverson57/campy). A side camera view was used for all analyses in this study. Tests of climbing behavior were performed during the animal’s dark cycle, illuminated by red and infrared lights.

Videos of climbing behavior were cropped to only the time when the animal is actually on the rod using either DeepEthogram^123^ or DeepLabCut^119^. Data was tracked in two dimensions using a DeepLabCut. All downstream analysis was performed using data that had a >0.65 likelihood of detection of the back, as reported by DeepLabCut. Data was smoothed using a 0.02-second Gaussian kernel. Time upright was calculated by calculating the difference between the minimum and maximum positions of the back. If it was greater than 100 pixels, the animal was determined to have flipped upside down, and the midpoint between the minimum and maximum value was considered to be the cutoff for calling upright or upside down. Most animals spent very little time near that midpoint value. Whether or not a mouse was moving was determined by a velocity threshold of greater than 5 pixels/frame.

### Generation and analysis of F2 hybrids

Four pairs of forest and prairie cross-subspecies mating pairs were set up in both directions (i.e., with the female being forest or prairie). The progeny of those crosses (F1s) were then crossed to animals that were not their siblings to generate F2 animals. 51 animals were run on the behavior assay. To facilitate performing the experiment on all animals, we ran them on three of the rod sizes: 12.7mm, 6.35mm, and 3mm. A subset of animals (5) was not run on the 6.35mm rod. Video data was missing for one day for two animals due to errors in video recording. Weights were measured at the time of the behavior. Foot, tail, and body length were measured at the time of sacrifice, and only 41 animals had foot, tail, and body length measurements taken due to ongoing behavioral experimentation outside of the scope of this study. To choose animals to use for the CST size analysis, we set thresholds for the fraction of time they spent upright and their velocities, which are not correlated with one another. We determined “good” climbers to be in the better half of all mice on both metrics and “bad” climbers to be in the worse half. Eleven mice were classified as “good” climbers, and eleven as “bad” climbers, and we picked randomly among them. We then watched videos to qualitatively confirm good and bad climbers. These animals were transcardially perfused, and spinal cords were stained as described above.

## Acknowledgements

K.M.T. was funded by a Life Sciences Research Foundation Postdoctoral Fellowship sponsored by the Howard Hughes Medical Institute (HHMI) and a NIH K99 BRAIN Initiative Advanced Postdoctoral Career Transition Award (K99NS133031). K.E.C. and D.C.E. were supported by the Harvard College Research Program and the Program for Research in Science and Engineering. This work was supported by laboratory startup funds (A.W.H.) and the HHMI (H.E.H). The Harvard Center for Biological Imaging assisted with light microscopy; the Harvard Center for Brain Sciences Neurotechnology core assisted with behavioral assay set-up and engineering; the Harvard Medical School Electron Microscopy Core assisted with electron microscopy; the Bauer Core at Harvard performed sequencing; the Image Analysis Collaborative at Harvard Medical School provided advice on whole-brain retrograde tracing analysis. We thank Wei-Chung Lee for advice on electron microscopy protocols; Caroline Hu for early work on viral tracing in deer mice and advice on surgeries; Chris Kirby and the Harvard Office of Animal Resources for animal husbandry; Jenny Chen for advice on single-cell data analysis and assistance with single-cell library generation; Kevin Cross and Stefen Lemke for advice on neural activity data analysis; Nick Tustison for advice on whole brain registration; and members of the Hoekstra lab for helpful discussion throughout the project. Evan Kingsley, Andi Kautt, Nacho Sanguinetti, Abhilasha Joshi, and Jenny Chen provided feedback on the manuscript.

## Data Availability

Data will be deposited in public databases upon publication. Data are available upon request.

## Author Contributions

K.M.T.: Conceptualization, Data Curation, Formal Analysis, Funding Acquisition, Investigation, Methodology, Project Administration, Software, Supervision, Visualization, Writing - Original Draft Preparation, Writing - Review & Editing; J.D.C.: Data Curation, Investigation, Methodology, Writing - Review & Editing; J-Z.G.: Data Curation, Investigation, Methodology; P.R.R.: Data Curation, Investigation, Methodology; K.E.C.: Data Curation, Investigation, Methodology; I.H.S: Data Curation, Investigation; D.C.E.: Data Curation, Investigation; A.W.H.: Conceptualization, Funding Acquisition, Supervision, Writing - Review & Editing; H.E.H.: Conceptualization, Funding Acquisition, Supervision, Writing - Review & Editing

## Notes

### Competing Interest Statement

The authors have declared no competing interest.

### Summary of Updates

Add additional neurophysiological analysis (Supplementary Figure 5 F-H). Add supplemental videos. Minor revisions to the text, most notably to the discussion section.

